# Co-movement of astral microtubules, organelles and F-actin suggests aster positioning by surface forces in frog eggs

**DOI:** 10.1101/2020.06.17.154260

**Authors:** James Pelletier, Christine Field, Sebastian Fürthauer, Matthew Sonnett, Timothy Mitchison

## Abstract

How bulk cytoplasm generates forces to separate post-anaphase microtubule (MT) asters in *Xenopus laevis* and other large eggs remains unclear. Previous models proposed dynein-based organelle transport generates length-dependent forces on astral MTs that pull centrosomes through the cytoplasm, away from the midplane. In *Xenopus* egg extracts, we co-imaged MTs, endoplasmic reticulum (ER), mitochondria, acidic organelles, F-actin, keratin, and fluorescein in moving and stationary asters. In asters that were moving in response to dynein and actomyosin forces, we observed that all cytoplasmic components moved together, i.e., as a continuum. Dynein-mediated organelle transport was restricted by interior MTs and F-actin. Organelles exhibited a burst of dynein-dependent inward movement at the growing aster surface, then mostly halted inside the aster. Dynein-coated beads were slowed by F-actin, but in contrast to organelles, beads did not halt inside asters. These observations call for new models of aster positioning based on surface forces and internal stresses.

## Introduction

Cytokinesis requires drastic reorganization of the cell, and it provides a model for probing cytoplasmic mechanics and principles of sub-cellular organization. Here, we focus on organization of the cytoplasm by MT asters that establish the cleavage plane before furrow ingression in *Xenopus laevis* eggs. Due to their ~1 mm diameter, these cells provide a system where organization of bulk cytoplasm by MT asters is particularly clear. The first mitotic spindle is centrally located, and much smaller than the egg. After mitosis, a pair of MT asters grow out from the centrosomes, reaching the cortex ~20 min later. These asters are composed of a branched network of short, dynamic MTs ~15 μm long and oriented approximately radially, with plus ends outward (Ishihara et al., 2016; Ishihara et al., 2014).

Asters have two important organizational and mechanical functions. The first function is to position cleavage furrows. Where the paired asters meet, at the midzone of the egg, the MTs form an antiparallel interaction zone which recruits the chromosomal passenger complex (CPC) and centralspindlin (Field et al., 2015), which define the cleavage plane (Basant et al., 2018; Carmena et al., 2012). The second function of MT asters, and the focus of this paper, is to move centrosomes and nuclei away from the future cleavage plane, so each daughter blastomere inherits one of each. This separation movement transports centrosomes and nuclei hundreds of microns away from the midplane over tens of minutes. Aster growth and centrosome separation movement after anaphase of first mitosis are illustrated in Figs 1A-C. In common with other authors, we often refer to centrosome and aster movement together for simplicity. The reality is more complex due to continuous MT growth and turnover. As centrosomes move away from the midplane, the aster surface adjacent to the midplane remains stationary, while the outer aster surface grows outwards due to a combination of MT polymerization and outward sliding. If the aster were rigid, surface MTs and centrosomes would slide outward at the same rate, but since the aster is better considered a deformable network, this is not a safe assumption.

**Figure 1.**
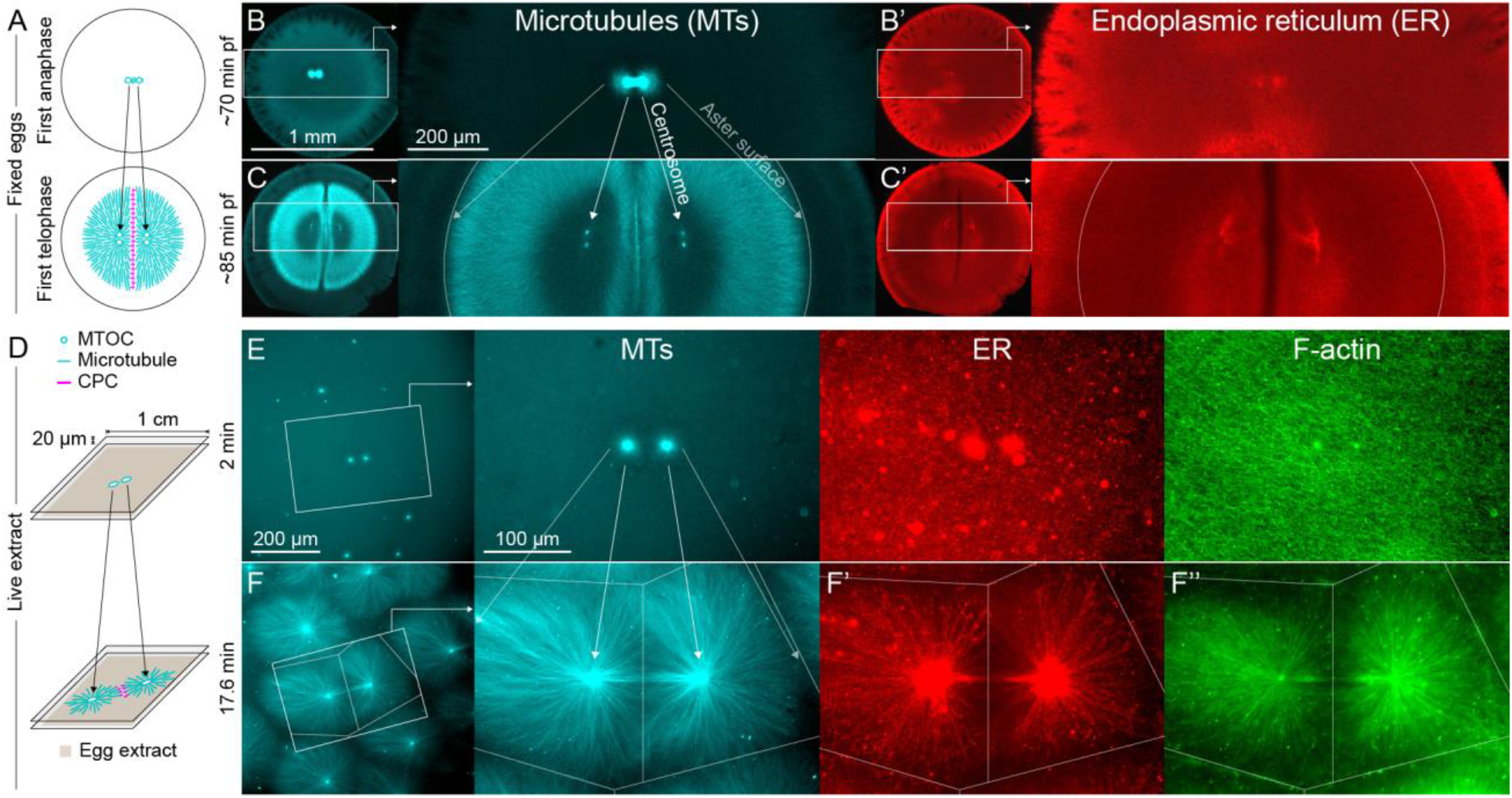
MTOC separation movement in *Xenopus* eggs and egg extract. Panels A-C are fixed embryos, and panels D-F are in *Xenopus* egg extract. (A) Cartoon illustrating MTOC movement away from the CPC-positive midplane before astral microtubules (MTs) reach the cortex in *Xenopus laevis* eggs. MTs shown in cyan and CPC-positive interaction zone in magenta. Note the CPC is shown in the cartoon panels A and D, but not in the rest of the figure. (B,C) Anti-tubulin immunofluorescence of eggs fixed ~70 and ~85 min post fertilization (pf). Diagonal lines connecting different eggs in panels B and C emphasize centrosome separation movement and growing aster surfaces. (B’,C’) Anti-LNPK (ER) immunofluorescence of the same eggs. (D) Cartoon illustrating aster separation movement in an extract system. MTs and CPC as in panel A. Asters were reconstituted from artificial microtubule organizing centers (MTOCs) in interphase *Xenopus* egg extracts. (E,F) MTOCs moved apart as asters grew and interacted with one another over time. Time is defined with respect to perfusing the sample and warming to 20 °C, so the start of aster growth occurred soon after 0 min. (F’) A fraction of the ER became enriched around MTOCs, and (F’’) F-actin was disassembled locally along interaction zones.

The forces that act on MTs to move centrosomes and asters have been extensively investigated and reviewed, and differ between systems (Garzon-Coral et al., 2016; Grill et al., 2005; Kotak et al., 2013; Meaders et al., 2020; Xie et al., 2020). In *Xenopus* and zebrafish zygotes, which are unusually large cells, Wühr and colleagues showed that movement away from the midplane after first mitosis is driven by dynein-dependent pulling forces (Wühr et al., 2010). Since movement occurs before astral MTs reach the cortex, the dynein must be localized throughout the cytoplasm, presumably attached to organelles, but this was not tested. The most prominent model for centrosome movement of this kind proposes that dynein attached to organelles generates pulling forces that increase with MT length, which can potentially account for the force asymmetries required for directed movement (Hamaguchi et al., 1986; Kimura et al., 2011; Tanimoto et al., 2016; Tanimoto et al., 2018; Wühr et al., 2010). In this “length-dependent pulling” model, dynein transports organelles along astral MTs toward the centrosome, then viscous or elastic drag on the organelles imparts a counter force on the MTs, pulling them away from the centrosome. The flux of organelles, and thus the net pulling force, is thought to scale with MT length. Although length-dependent pulling models are widely discussed, many aspects remain unclear. Net forces may not scale with MT length due to hydrodynamic interactions between MTs, which computational studies have shown are significant (Nazockdast et al., 2017). The organelles that anchor dynein in the cytoplasm of large egg cells have not been fully identified and the spatiotemporal distribution of organelle transport has not been mapped. Candidate dynein anchor organelles include the ER, which moves inwards as sperm asters center in sea urchin (Terasaki et al., 1991), acidic organelles which were implicated in nematode eggs (Kimura et al., 2011), and mitochondria which are abundant in early embryos.

Contractile activity of actomyosin can cause centrosome and aster movement in eggs and embryos (Field & Lenart, 2011; Telley et al., 2012), but its contribution to centrosome separation movement in *Xenopus* eggs is unclear. Bulk cytoplasmic F-actin is a major mechanical element in *Xenopus* eggs (Elinson, 1983) and egg extracts (Field, Wühr, et al., 2011). In egg extracts, F-actin can impede centrosome movement in meiotic extracts (Colin et al., 2018), but F-actin is not required for centrosome separation movement in cycling extracts (Cheng et al., 2019). A note of caution is required in interpreting drug studies. F-actin depolymerization softens the cytoplasm, and presumably decreases the drag on moving asters as well as dynein anchors. Thus, F-actin depolymerization may modulate dynein-based forces on asters, in addition to removing actomyosin-based forces. Furthermore, effects of cytochalasins in *Xenopus* eggs are hard to interpret because they only permeate the *Xenopus* egg cortex during first cleavage, when new membrane becomes exposed (de Laat et al., 1973).

Considering both the length-dependent pulling model, and the role of actomyosin, one important question has not been asked in *Xenopus* eggs: do centrosomes and associated astral MTs move *through* the cytoplasm and the other networks, as predicted by the length-dependent pulling model? Or do they move *with* the other networks as a continuum? If organelles anchor dynein, the length-dependent pulling model predicts that the organelles must move in the opposite direction as the centrosome, or at least remain stationary. Live observation is required to compare movement of MTs and organelles. This is not possible in opaque frog eggs so we turned to actin-intact egg extracts. Growth and interaction of interphase asters were previously reported in this system (Ishihara et al., 2014; Nguyen et al., 2014), but aster movement was not systematically investigated. Here, we report methods for observing aster movement in egg extract and use them to measure relative movement of MTs, organelles, and F-actin. We observed all cytoplasmic networks moved together as a continuum inside asters, whereas the highest velocity differences between networks occurred at aster surfaces. We end by proposing a new “surface force” model for positioning centrosomes and asters in eggs.

## Results

### Centrosome separation and ER distribution in fixed eggs

As a first test of how centrosomes and organelles move relative to one another, we fixed frog eggs before first cleavage, stained for tubulin and ER, and imaged by confocal microscopy (Figs 1A-C). Centrosome separation movement is represented by the diagonal lines connecting different eggs in panels B and C. As centrosomes move away from the midplane, the centrioles within them replicate and split, visible as the pair of bright cyan spots within each aster in Fig 1C. We probed the ER distribution by staining for the ER membrane marker Lunapark (LNPK) (Figs 1B’,C’) and also the luminal marker PDIA3 (not shown) with similar results. The ER was distributed all over the asters, with some enrichment near centrosomes and the cortex. Lack of strong ER enrichment at centrosomes called into question the length-dependent pulling model with the ER as a dynein anchor. However, organelle transport dynamics could not be measured from fixed images, so we turned to an egg extract system for live imaging.

### Microtubule organizing center (MTOC) separation movement in egg extract by dynein and actomyosin

To model separation movement in a cell free system suitable for live imaging, we filled chambers consisting of two PEG-passivated coverslips spaced ~20 μm apart with actin-intact interphase egg extract containing artificial MTOCs, imaging probes, and drugs. MTOCs nucleated astral MTs. We then imaged aster growth and MTOC movement over ~30 min. For most experiments we used widefield microscopy with a 20x objective to collect data on overall organization and flows, in some cases stitching multiple image fields. To illustrate structural details of the components we studied, Fig S1 and SI Movie 1 show MTs, ER, and F-actin near an MTOC by spinning disk confocal microscopy with a 60x objective. In 20x magnification fields, we routinely noted that MTOCs that were close together at early time points tended to move apart. Fig 1D illustrates the extract system, and Figs 1E,F show an example of MTOCs moving apart as asters grew and interacted with one another. This kind of separation movement was observed in hundreds of image sequences, such as in SI Movie 2, and we believe it models centrosome separation movement in eggs.

When asters grew to touch each other, they formed CPC-positive interaction zones (Fig 2A) as previously reported (Nguyen et al., 2014). These reconstitute the CPC-positive zone that forms between asters in eggs (Field et al., 2015). CPC-positive zones cause local disassembly of both MTs and F-actin, which locally softens the cytoplasm (Field et al., 2019), and generates strong anisotropy in both MT and F-actin density. These anisotropies may lead to generation of directed forces on MTOCs by both dynein and actomyosin. Previous work focused on possible consequences of MT length anisotropy on dynein forces (see Introduction). Here, we will focus on consequences of locally softening the cytoplasm at CPC-positive zones on the mechanical response of asters to forces from both dynein and actomyosin.

**Figure 2.**
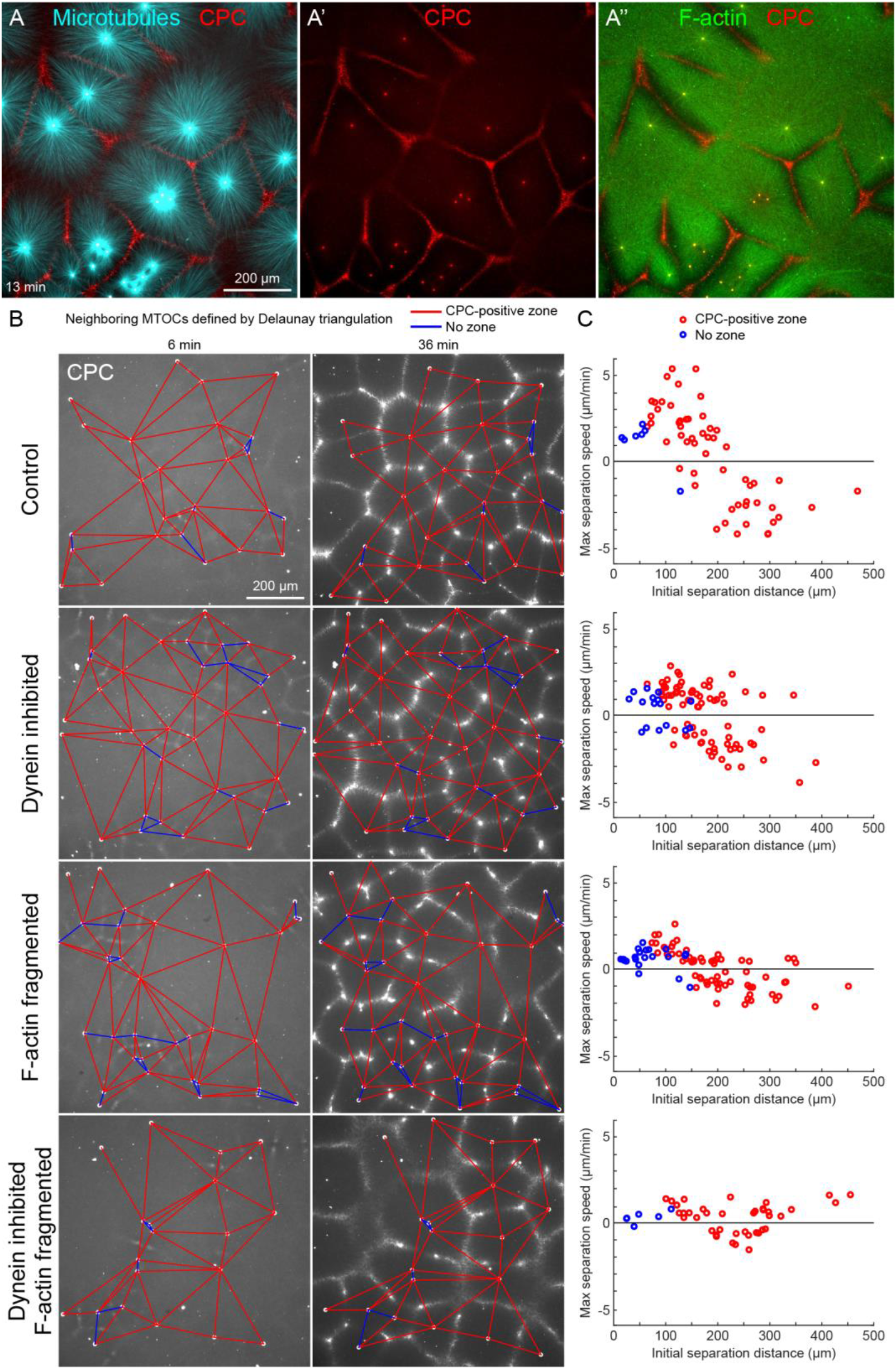
MTOC separation movement in egg extract by dynein and actomyosin. (A) The CPC localized to interaction zones between neighboring asters, blocking mutual interpenetration of MTs and disassembling F-actin locally. Time is defined with respect to perfusing the sample and warming to 20 °C, so the start of aster growth occurred soon after 0 min. (B) Four aster growth reactions were followed in parallel under control vs inhibitor conditions. The first column in each condition shows an early time point, and the second column shows a time point 30 min later. MT growth was similar and CPC-positive zones formed under all conditions (see SI Movie 3). (C) Maximum speed of separation with respect to initial distance between the MTOCs.

To quantify MTOC movement, and determine the role of forces from different motors, we picked random locations and imaged large fields over time in up to four conditions in parallel. Fig 2B and SI Movie 3 show a typical experiment, where only the CPC channel is shown for simplicity. At early times points, the spatial distribution of MTOCs was random and the CPC signal was diffuse, except some signal on the MTOCs. As asters grew and interacted, they recruited CPC to zones between them under all conditions. We quantified MTOC movements with respect to their nearest neighbors, which were defined by the Delaunay triangulation between MTOCs at the earliest time point and followed over the movie (Fig 2B). Red edges indicate when neighboring MTOCs formed a CPC-positive zone between them, and blue edges indicate when they did not. We then measured the maximum separation speed as a function of the initial separation distance between the MTOCs.

Under control conditions, MTOCs that were initially closer together tended to move farther apart, while those initially farther apart tended to move closer together, as evidenced by the strong negative correlation between the maximum speed of separation movement and starting distance (Fig 2C). We focused on separation movement of MTOCs in separate asters with a CPC-positive zone between them (red points), since this models centrosome separation movement in eggs.

To test the role of dynein and actomyosin in MTOC movement, we inhibited dynein or fragmented F-actin, separately or together. Inhibiting only one motile system caused a partial block. The role of each motile system was similar, as judged by effects on the slopes of separation speed vs initial distance plots (Fig 2C). We investigate sites of dynein-based pulling below. We hypothesize actomyosin-based separation movement is driven by actomyosin contraction away from regions of lower F-actin density along interaction zones, and will analyze this model in detail elsewhere. Inhibiting both motile systems completely blocked MTOC movement. Inhibiting CPC recruitment with an AURKB inhibitor also completely blocked MTOC movement (not shown). These findings were qualitatively confirmed by visual inspection and partial analysis of more than 10 experiments using multiple extracts. We interpret these data as showing that MTOC movement in our extract system is driven by a combination of dynein and actomyosin forces. We next focused on analysis of co-movement of astral MTs, organelles and F-actin during aster separation movement.

### ER and F-actin move with MTs in separating asters

MTOC separation trajectories were longest, and most unidirectional, when MTOCs were clustered at the initial time point. In these cases, MTOCs moved predictably outwards from the cluster as asters grew out and interacted (Figs 3A,B, SI Movie 2). In Fig 3A, future MTOC trajectories are superimposed on an early time point to illustrate separation movement. To investigate how ER and F-actin moved with respect to moving astral MTs, we first used kymograph analysis. We picked a pair of MTOCs that moved apart, indicated by stars in Figs 3A-C. Then we generated kymographs in all channels (Fig 3C) along the line passing through the MTOCs, indicated by the grey line in Figs 3A,B. Visual inspection revealed features in all three channels that tracked parallel to the separating MTOCs, suggesting all the networks were moving together away from the interaction zone, on both the leading and trailing sides of the aster indicated in Fig 3B. Organelles visible in differential interference contrast (DIC) images also moved away from the interaction zone (SI Movie 2). These features are most evident in the F-actin kymograph, but can be seen in all channels by magnifying the figure and inspecting closely. Visual inspection and kymograph analysis of image sequences from more than 10 independent experiments confirmed that all components of asters tend to move together during separation movement, and that the data in Fig 3 are typical.

**Figure 3.**
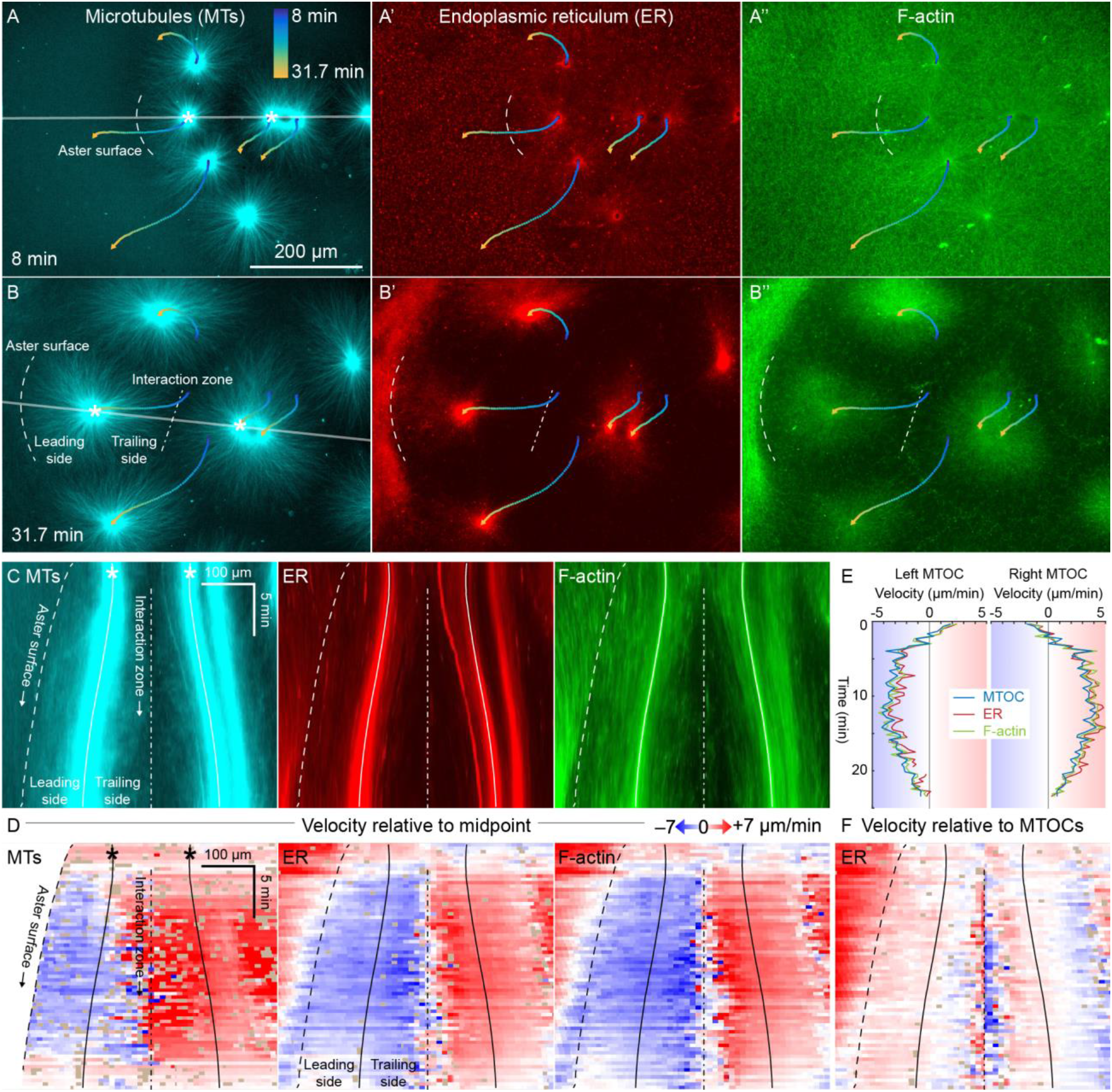
ER and F-actin move with MTs in separating asters. (A,B) Asters grew until they reached their neighbors, formed interaction zones approximately equidistant between the MTOCs, then moved away from the interaction zones (see SI Movie 2). MTOC trajectories are represented by contours colored from blue to yellow. Time is defined with respect to perfusing the sample and warming to 20 °C, so the start of aster growth occurred soon after 0 min. (C) Intensity kymographs along the grey line shown in panels A and B, passing through the MTOCs marked with a white star. To show relative movement of the MTOCs, each row of the kymograph was computationally translated to keep stationary the midpoint between the MTOCs, where the interaction zone formed. Solid curves indicate the MTOCs, the dashed curve indicates the growing aster surface, and the dash-dotted line indicates the interaction zone. (D) Velocity kymographs in the same frame of reference as in panel C. 2D flow fields were measured by particle image velocimetry (PIV), projected onto the line passing through the MTOCs, then the projected velocity of the midpoint between the MTOCs was subtracted, again to show movement relative to the interaction zone. A white color indicates stationary with respect to the midpoint, blue indicates moving to the left, and red to the right. PIV outliers were filtered and shown in beige. (E) Velocity of the MTOCs based on particle tracking, as well as the velocity of ER and F-actin in the neighborhood of the MTOCs based on PIV. (F) Velocity of ER with respect to the moving MTOCs, not with respect to the interaction zone as in panel D.

To quantify how MTs, ER, and F-actin flowed relative to one another during aster movement, their 2D flow fields were independently measured by particle image velocimetry (PIV). All three cytoplasmic networks moved in the same direction at similar speeds of up to 7 μm/min. Movement was always directed away from the interaction zone on both leading and trailing sides of the aster, as shown by blue on both sides annotated in Figs 3C,D. This is inconsistent with the length-dependent pulling model, in which organelles on the leading side must move toward the interaction zone (red) or at least remain stationary with respect to the interaction zone (white). Fig 3E compares the MTOC velocity from particle tracking to the ER and F-actin velocities from PIV, again consistent with all three cytoplasmic networks moving at similar speeds. To highlight relative movement within asters, Fig 3F shows the ER velocity relative to the MTOC velocity. Relative movement within asters was ~20% the speed of all networks away from the interaction zone, as evidenced by the pale colors in Fig 3F. In contrast, the highest velocity gradients were observed near the aster surfaces (Figs 3D,F).

Close inspection of Fig 3D and similar analyses showed that velocities of all three networks away from the interaction zone were not constant throughout the aster, though different networks had similar velocities at any given location. Typically, the region near the interaction zones moved ~20% faster than the MTOC, and the leading edge of each aster moved ~20% slower. These PIV-based velocities reflect polymer translocation rather than polymerization, so this spatial variation in velocity shows that the aster does not move as a completely rigid body. Rather, it deforms as a gel, compressing or stretching in response to forces and stresses. Disassembly of F-actin softens the interaction zone, facilitating aster separation movement.

### ER and F-actin move with MTs on coverslips functionalized with dynein

To generate a complementary system for dynein-dependent MTOC movement where actomyosin was less important, we artificially anchored dynein to the coverslip via a biologically relevant linkage. This also had the effect of speeding up movement ~10 fold compared to movement away from CPC-positive zones. Endogenous HOOK2, a coiled-coil dynein-dynactin adapter (Reck-Peterson et al., 2018), was recruited to PEG-passivated coverslips via an antibody raised to its C-terminus (Methods). To characterize the antibody and identify HOOK2 interacting proteins, we performed quantitative immunoprecipitation-mass spectrometry (IP-MS) (Fig S2). We compared 3 conditions: anti-HOOK2 in interphase extracts (3 separate extract repeats), anti-HOOK2 in mitotic extracts (2 repeats), and as negative control, random IgG in interphase extracts (3 repeats). HOOK2 was the most abundant protein recovered on anti-HOOK2 beads. HOOK3 was also detected, consistent with heterodimerization between HOOK family members (Redwine et al., 2017; Xu et al., 2008). In interphase extracts, anti-HOOK2 pulled down multiple subunits of the dynein-dynactin complex, plus known interactors LIS1 and CLIP1. All these dynein-related proteins were greatly reduced in pulldowns from mitotic extracts, suggesting the interaction between HOOK2 and dynein-dynactin is negatively regulated by CDK1 activity. We also noted many potential dynein-dynactin interacting proteins that have not been confirmed by other methods (data available on request). We concluded the HOOK2 antibody offers a physiological linkage to dynein, and we proceeded to test its effects on aster movement.

Dynein attached to coverslips via HOOK2 generated pulling forces on MTs directed away from the MTOC (Fig 4A). We previously reported that dynein non-specifically adsorbed to non-passivated coverslips increases the rate of aster growth due to outwards microtubule sliding, but did not move MTOCs (Ishihara et al., 2014). Remarkably, on HOOK2-functionalized coverslips, asters exhibited rapid translational movement in a circular pattern with a diameter of 20-30 μm (Figs 4B,C, SI Movie 4). During this movement, MTOCs moved continuously at ~1 μm/s, comparable to the maximum speed of dynein (Reck-Peterson et al., 2018) (Fig 4D). This 2D-oscillatory movement was observed in >10 different experiments using different batches of extract, and was blocked by dynein inhibition. We plan to investigate the mechanism of the instability that causes circular motion elsewhere. Here, we used the rapid aster movement as an alternative system to study how ER and F-actin move with respect to moving MTs. Fig 4E shows intensity kymographs along a horizontal line that tracks up and down with the MTOC, analogous to the kymographs in Fig 3C. Fig 4F shows velocity maps analogous to those in Fig 3D. The intensity kymographs reveal many features that tracked with the MTOC, and the velocity plots show that indeed, all the cytoskeletal networks moved in the same direction, at the same speed, at any location inside the aster. As with separation movement in Fig 3, the asters gliding over HOOK2-functionalized coverslips moved as a continuum. In another experiment, keratin was also advected with moving asters (SI Movie 5). From these observations we conclude that cytoplasmic networks are mechanically integrated inside asters, and cytoplasmic networks move together with moving asters.

**Figure 4.**
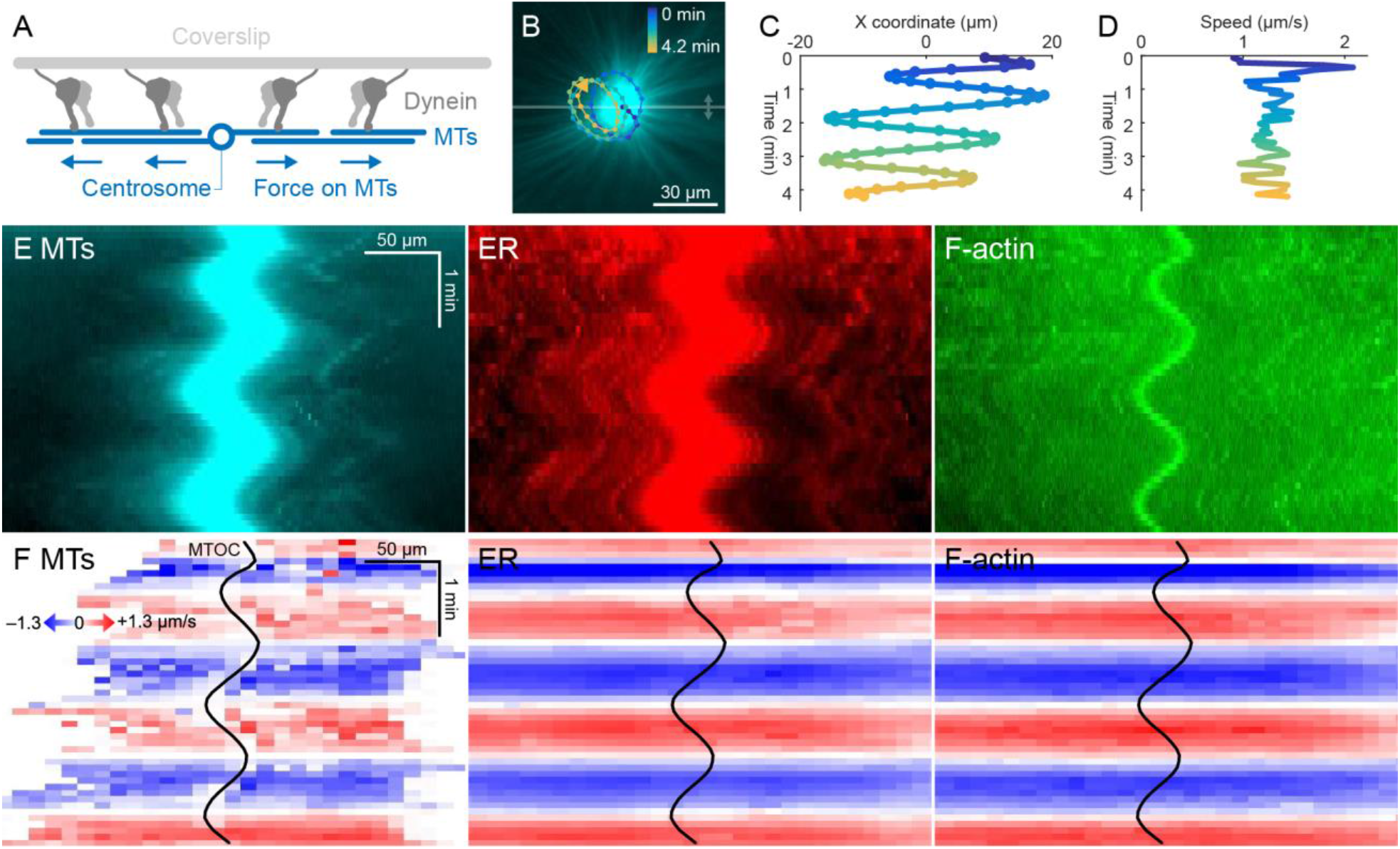
ER and F-actin move with MTs on coverslips functionalized with dynein. (A) Coverslips were functionalized with an antibody against HOOK2, so the rigid coverslip substrate generated pulling forces on the astral MTs. (B) Circular oscillatory trajectory of the MTOC (see SI Movie 4). (C) X coordinate of the MTOC. (D) Speed of the MTOC relative to the coverslip, including both X and Y components of motion. (E) Intensity kymographs along the horizontal line passing through the MTOC, indicated in panel B. (F) Velocity maps in the same frame of reference as in panel E. 2D velocity fields were measured by particle image velocimetry (PIV) then projected onto the horizontal line as in panel E. The MTOC position is shown as a black curve.

### A small molecule probe is advected with moving asters

The high speed and predictability of the oscillatory movement on HOOK2-functionalized coverslips enabled us to ask whether the cytosol was advected with the moving cytoplasmic networks. This question was inspired by recent experiments showing that moving actomyosin gels advect cytosol in *Drosophila* embryos (Deneke et al., 2019). We functionalized artificial MTOCs with caged fluorescein, linked to the MTOCs via the caging group (Fig 5A). The fluorescein was uncaged upon shining 395 nm light, simultaneously activating its fluorescence and releasing it from the MTOCs (Figs 5A-C, SI Movie 6). The cloud of photo-released fluorescein dispersed within tens of seconds (Fig 5D). Rapid diffusive spread of the cloud validated that the fluorescein behaves as a freely diffusing small molecule (Fig 5E) and enabled estimation of the viscosity of the cytosol at ~6x that of water (Methods), consistent with previous estimates (Luby-Phelps, 1999; Valentine et al., 2005). We then fit the fluorescein cloud with a 2D Gaussian to track its center of mass. The center of brightness of the diffusing fluorescein cloud was clearly advected with the MTOC (Fig 5F), showing that cytosol advects with moving asters due to hydrodynamic interactions inside asters. Advection of cytosol with moving asters is consistent with poroelastic behavior and places an upper bound of ~100 nm on the effective pore size of cytoplasmic networks in this system (Methods) (Mitchison et al., 2008; Moeendarbary et al., 2013). Similar evidence of advection was found in >10 experiments in 3 extracts. The cloud center and the MTOC did not precisely co-align. This could be due to tracking error, but we suspect that bulk liquid flow driven by forces outside the aster may permeate asters and move the diffusing cloud.

**Figure 5.**
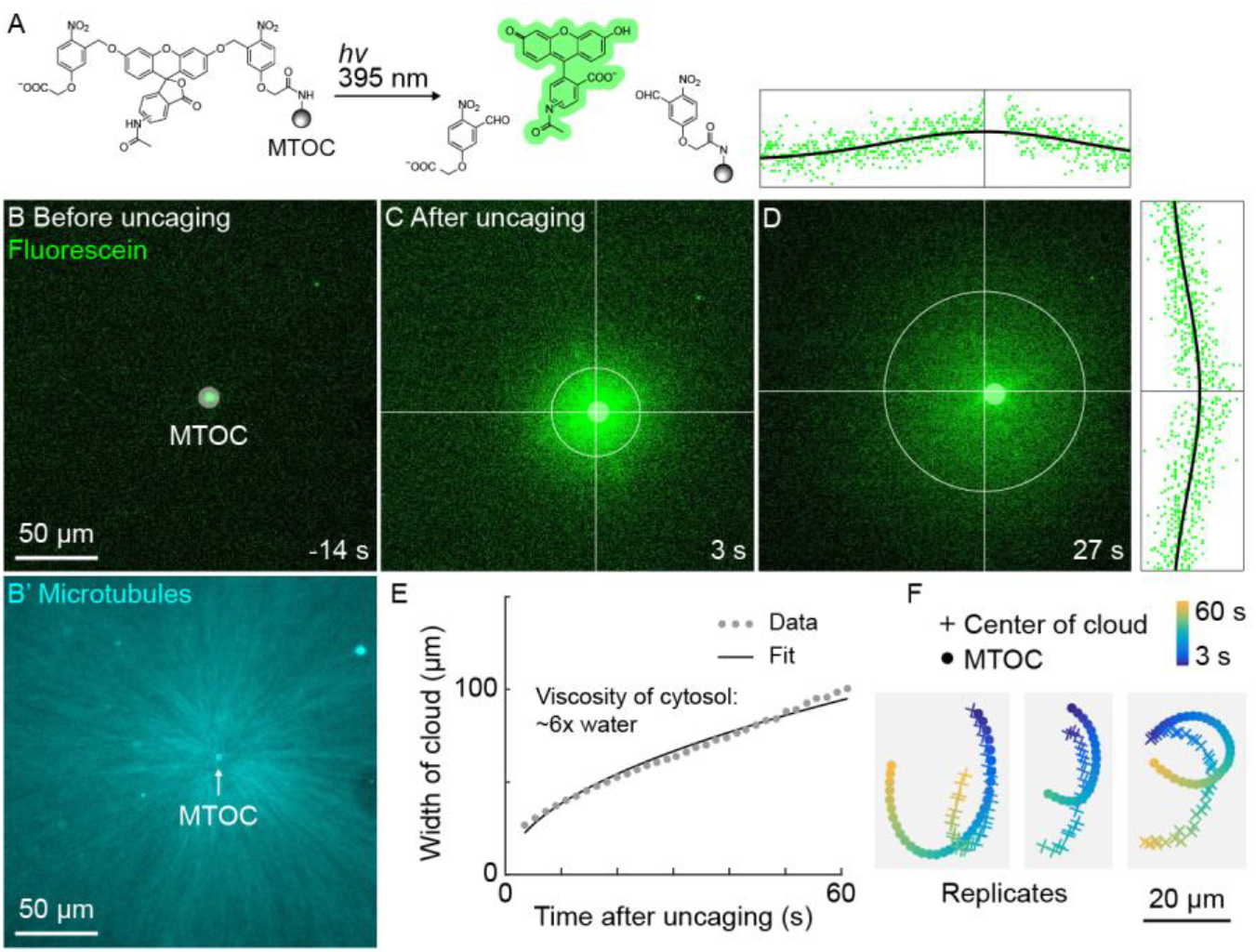
A small molecule is advected with moving asters. (A) To track the flow of a small molecule within moving asters, MTOCs were functionalized with caged fluorescein. (B) Caged fluorescein, before uncaging. (B’) Astral MTs radiating from the MTOC filled the region. The aster was oscillating on a coverslip functionalized with anti-HOOK2 as in Fig 4. (C) Fluorescein, after uncaging. (D) Within tens of seconds, the fluorescein diffused away from the MTOC and approached the background intensity (see SI Movie 6). 2D Gaussian fits to estimate the width and center of the fluorescein cloud. The bright MTOC was excluded from the Gaussian fit, so uncaged fluorescein that remained bound to the MTOC did not bias the fitted position. (E) Expansion of the fluorescein cloud width fit to a model of diffusion. (F) Several replicate trajectories of the center of the fluorescein cloud (plus) and the MTOC (circle).

### Dynein-mediated organelle movement is restricted by F-actin and interior MTs

We next investigated which organelles in egg extract recruit dynein, and how they might exert force on asters in a continuum model for aster movement. To facilitate detailed analysis of organelle transport throughout the aster, we imaged isolated asters which remained stationary as they grew. ER and mitochondria are the most abundant organelles in *Xenopus* egg extracts based on proteomics (Wühr et al., 2014), and acidic organelles were implicated in centrosome movement in *C. elegans* embryos (Kimura et al., 2011).

In control extracts with F-actin intact, almost all the ER, mitochondria, and acidic organelles remained evenly distributed over asters as they grew, and a small fraction of the ER accumulated near MTOCs (Figs 6A, 7A,D, SI Movies 7,8). The ER intensity around MTOCs increased to ~2 fold higher than the intensity outside the aster (Fig 6A’) in >5 examples scored. Although the majority of ER remained stationary, astral MTs did induce a subtle change in the texture of the ER, from coarser outside the aster, to finer and more tubular in appearance inside the aster (SI Movies 7,8). Astral MTs also affected the structure of the F-actin network, from random orientation of filaments outside the aster, to radial alignment of a subpopulation of bundles inside the aster (Fig 7A) as we reported previously (Field et al., 2019).

**Figure 6.**
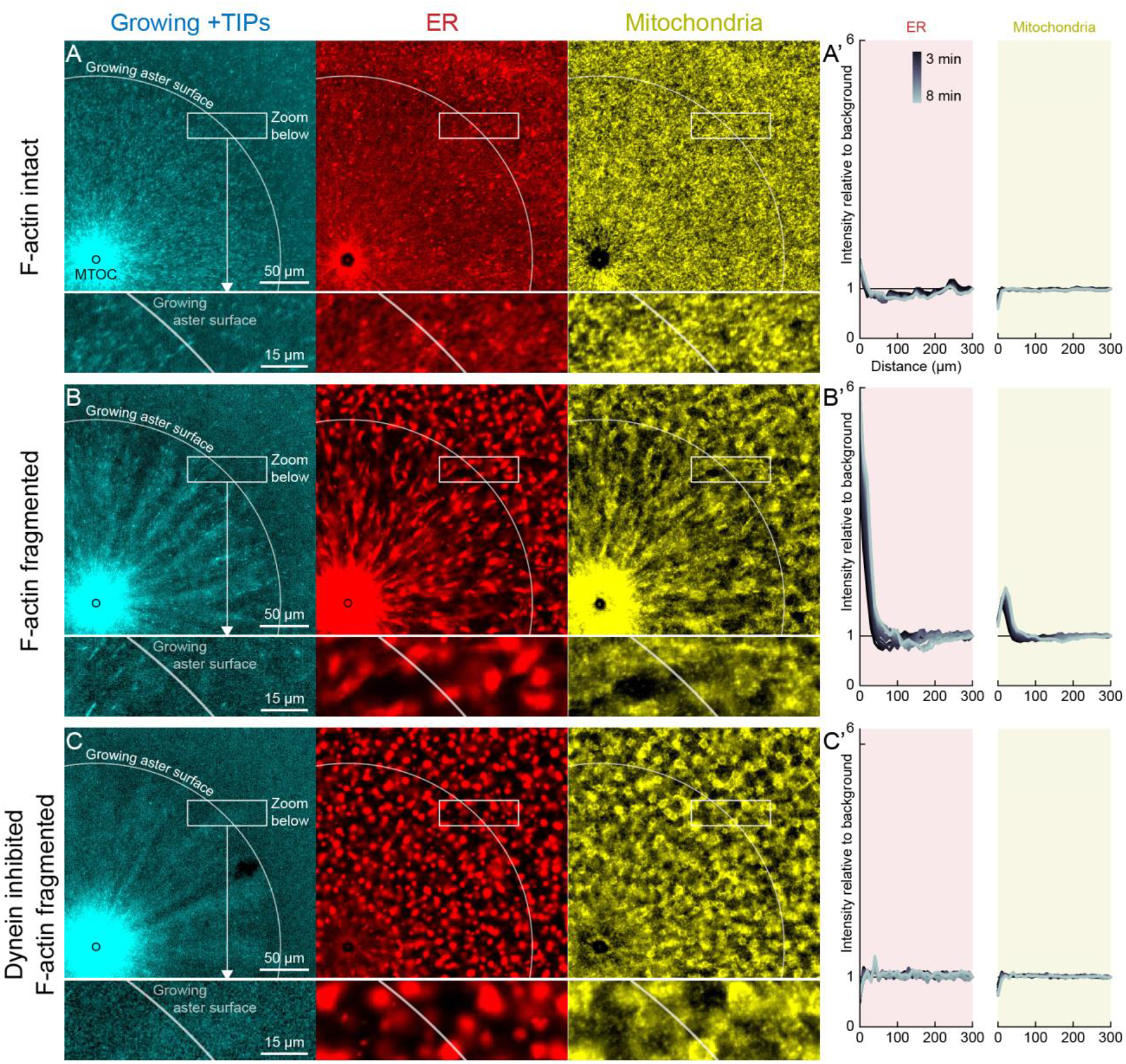
Dynein-mediated organelle movement is restricted by F-actin. (A) In control with intact F-actin, a small amount of ER became concentrated around the MTOC, but the majority of the ER and mitochondria remained distributed over the aster (see SI Movie 7). The white arc indicates the growing aster surface, and the box indicates the zoomed region in the lower panels. (A’) Average intensity with respect to distance from the MTOC over time, from black to gray. (B) When F-actin was fragmented with Cytochalasin D, a greater fraction of the ER was transported toward the MTOC, and a fraction of mitochondria was transported as well. Higher magnification: ER started to move when MTs indicated by growing +TIPs first grew into the cytoplasm, and ER and mitochondria co-localized with one another. (C) When dynein was inhibited with CC1, the ER was not transported, neither toward nor away from the MTOC.

**Figure 7.**
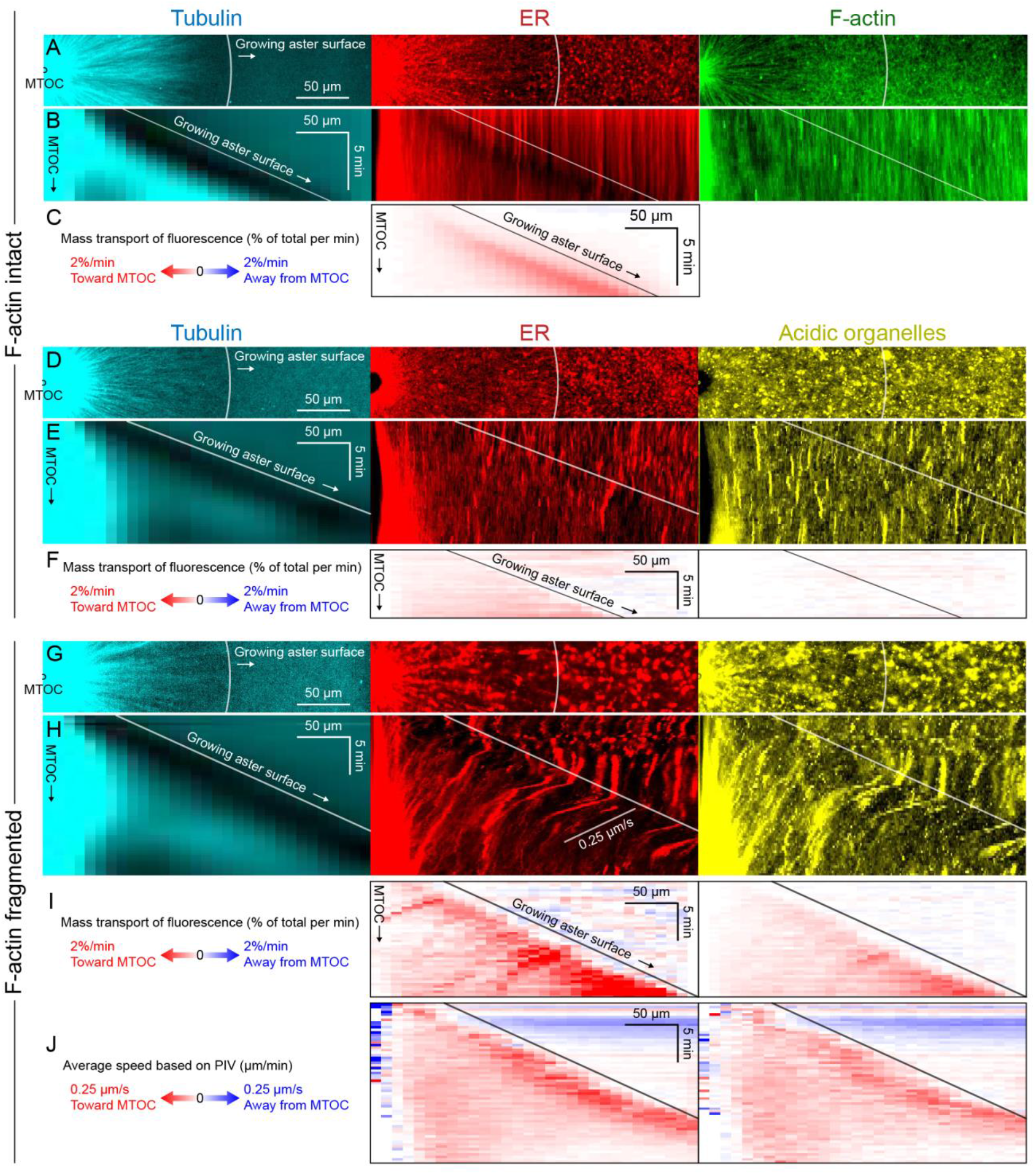
Dynein-mediated organelle movement is maximal on the aster surface. (A) Stationary asters were grown from isolated MTOCs. The growing aster surface is indicated by a white arc, and the ER was largely distributed but slightly depleted just inside the growing aster surface. The ER exhibited a change in texture from slightly coarser outside the aster to finer inside the aster (see SI Movie 8). (B) Kymographs along a line extending away from the MTOC. The MTOC corresponds to the left column, and the growing aster surface corresponds to the diagonal line where soluble tubulin is depleted upon incorporation into the growing aster. (C) Mass transport map for ER averaged over a quadrant, in the same frame of reference as the kymographs in panel B. Mass transport analysis is described in Fig S4. (D-F) Similar experiment with F-actin intact, in a different batch of extract that exhibited less organelle movement (see SI Movie 9). (G-J) Similar experiment with F-actin fragmented by Cytochalasin D (see SI Movie 10). (J) Average speed based on PIV, in the same frame of reference as panels H,I and averaged over a quadrant. PIV is not shown for control because movement was too slow to be reliably quantified.

When F-actin was fragmented with Cytochalasin D, all organelles exhibited inward movement (Figs 6B, 7G, Fig S3), which was fastest at the growing surface of the aster (Fig 7). Compared to control, a greater fraction of the ER was transported inwards (Fig 6B’), and average transport speeds were an order of magnitude faster with F-actin fragmented than intact (Fig 7). The ER intensity around MTOCs accumulated to ~6 fold higher than the intensity outside the aster and continued to increase with time (Fig 6B’). Due to the burst of movement at the surface of the growing aster, the intensity of organelles was ~30% lower there than outside the aster (Fig 6B’). Compared to control, the texture of the ER was coarser when F-actin was fragmented, both inside and outside asters, and MTs appeared more bundled. Mitochondria and acidic organelles moved inwards and accumulated near the MTOC. These organelles appeared to physically associate with ER in higher magnification images (Figs 6B,C, 7G, S3), so all organelles may be physically connected in this system. These findings show that the ER, and perhaps all organelles, recruit dynein, and can move toward the MTOC. Inward movement is restrained by F-actin under control conditions. However, even with F-actin fragmented, the majority of the ER, mitochondria, and acidic organelles were still evenly distributed over the aster.

We next added CC1 to test for a role of dynein in organelle transport (Fig 6C). We did this with and without Cytochalasin D to fragment F-actin, but only report results with F-actin fragmented because inward movement is much easier to score. With CC1 present, with or without F-actin, organelles moved neither inwards nor outwards, and did not accumulate at MTOCs. This suggests that dynein generates all the inward force on organelles, and that other forces on organelles do not induce significant net transport in this system. Tip-tracking and kinesin forces might escape detection in 20x movies if they only cause local movements, but they did not make a major contribution to overall organelle distributions.

### Dynein-mediated organelle movement is maximal near the aster surface

To infer outward forces on MTs as a function of time and location, we needed a measure of the total inward organelle flux. Kymographs and PIV provide direct visualization of movement but have limitations for this inference, because they measure movement of local gradients in fluorescence intensity, not mass transport of organelles. We therefore developed an analysis to infer mass transport of organelles based on flux of fluorescence intensity (analysis described in Fig S4 and Methods). This analysis quantifies the amount of fluorescence signal crossing a given circumference at a given time, normalized by the total fluorescence in a region containing the aster. Inward organelle transport from this analysis can be qualitatively related to outward force on MTs (Discussion). Fig 7 shows examples with ER and acidic organelles. Mitochondria exhibited similar movement as acidic organelles (Fig S3).

All analysis methods revealed a burst of inward organelle movement when the growing aster surface reached them, followed by slowing down inside asters (SI Movies 8-10). This burst can be visualized as inward diagonal features in kymographs, and high values on the surface diagonal in mass transport and PIV plots. Under control conditions, with F-actin intact, the amount of organelle movement at the aster surface was variable between extracts. Out of 11 extract preps, we observed a burst of inward ER movement at the aster surface in 7 extracts (64%) as in Fig 7C, and observed weaker or no burst in the remaining extracts as in Fig 7F. Factors that seem to lessen the burst of inward movement include higher concentrations of spontaneously nucleated MTs outside the aster, and insufficient passivation of the coverslips, but these two factors do not explain all the examples that did not exhibit a burst. Lack of fast organelle movement in control asters with intact F-actin is consistent with co-movement of cytoplasmic networks in moving asters (Figs 3 and 4).

When F-actin was fragmented to highlight interactions between organelles and MTs, organelle transport at the growing aster surface was faster, and therefore easier to visualize and quantify. The burst of inward movement near the aster surface was highly reproducible when F-actin was fragmented with Cytochalasin D. Velocity values for ER moving inwards at the aster surface reached ~0.25 μm/s with F-actin fragmented (Fig 7J), and mass transport reached 2% per min (Fig 7I). Mass transport values were more peaked at the surface than PIV values, in part because mass transport takes into account the increase in circumference as the radius increases. However, the PIV values were peaked at the surface, as well, so organelles moved faster near the surface then slower once incorporated into asters. The peak in velocity at the aster surface was highly reproducible with F-actin fragmented and observed in >10 experiments with separate extracts. A smaller fraction of acidic organelles than ER were transported inwards (Figs 7F,I), but with a similar bias toward movement at the surface. The restriction of organelle transport to the aster surface, even with F-actin fragmented, suggests that forces from dynein are exerted primarily at the growing aster surface.

### Dynein-coated beads move inwards at constant rates throughout asters

Slowing of organelle transport upon incorporation into the aster, whether F-actin was intact or fragmented, suggested dynein might be inhibited inside asters, for example by chemical signals. To test this, we turned to an artificial system. 3 μm beads were functionalized with the antibody against the dynein adapter HOOK2 used in Figs 4 and 5. Negative control beads were functionalized with random IgG. We then measured transport of the beads on isolated, stationary asters as in Figs 6 and 7. With F-actin intact, the anti-HOOK2 beads moved inwards at a constant speed of 0.2 ± 0.1 μm/s throughout asters (Figs 8A-D, SI Movie 11). When F-actin was fragmented with Cytochalasin D, the anti-HOOK2 beads moved at 0.7 ± 0.2 μm/s (Figs 8E-H), faster than with F-actin intact. Thus, artificial dynein-coated beads were slowed by F-actin, like endogenous organelles. However, these beads were transported all the way to the MTOC, unlike organelles which slowed or stopped inside asters. This suggests some brake on dynein movement is recruited to organelles but not to HOOK2-coated beads, or alternatively, that the force on HOOK2-functionalized beads is higher.

**Figure 8.**
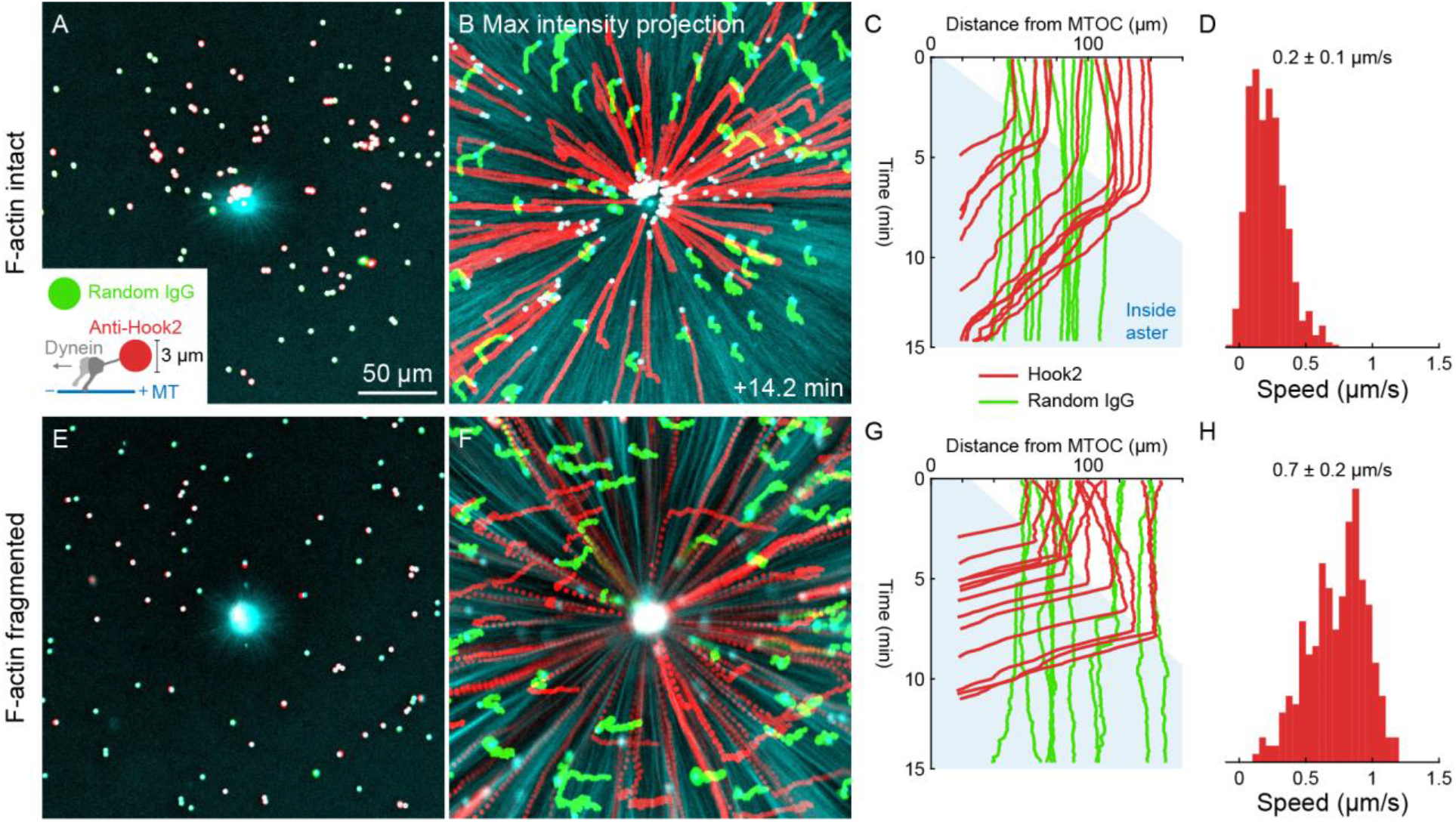
Unlike organelles, artificial cargoes functionalized with dynein move at constant speed throughout asters. (A) Artificial cargoes were functionalized with an antibody against the dynein adapter HOOK2, and negative control beads were functionalized with random antibody (see SI Movie 11). (B) Max intensity projections of beads functionalized with anti-HOOK2 (red) or random antibody (green). (C) Trajectories of anti-HOOK2 and negative control beads relative to the MTOC. The growing aster is indicated by blue region. Anti-HOOK2 beads started to be transported when they were engulfed by the growing aster, indicated by the blue region. (D) Velocity distribution of anti-HOOK2 beads inside the aster. (E-H) Similar experiment with F-actin fragmented by Cytochalasin D.

### Volume exclusion is unlikely to block organelle movement inside asters

Organelles might slow down inside asters because the environment becomes too crowded with other organelles. To investigate volume exclusion by organelles, we quantified the intensity and flux of fluorescent dextran as a marker for the cytosol. As organelles were transported toward MTOCs, fluorescent dextran was displaced away from MTOCs (Fig S5, SI Movie 12), consistent with volume conservation. However, the degree of steric exclusion was fairly small, since the dextran signal was only reduced by ~10%, and exclusion was only observed within ~50 μm of MTOCs, where the ER density is maximal. Outside that central region, the intensity of fluorescent dextran was similar inside and outside asters. We conclude that volume exclusion between organelles may be significant in the immediate neighborhood of MTOCs, but is unlikely to account for organelles becoming stationary inside asters.

## Discussion

We tracked multiple cytoplasmic networks in moving asters using two different systems to promote movement, and found that organelles, F-actin, keratin, and even a small molecule moved coherently with astral MTs (Figs 3,4,5, SI Movie 5). During aster separation movement, organelles on both the leading and trailing sides moved away from the interaction zone (Fig 3, SI Movie 2). In stationary asters, organelles exhibited a burst of movement at the aster surface, then mostly halted inside asters (Figs 6,7). This burst was much more pronounced when F-actin was fragmented, but even in that situation the majority of organelles were slower or stationary once they joined the body of the growing aster. Thus, asters moved as a near-continuum, with maximal shear between networks at the aster surface, and much less shear between networks inside asters. Co-movement of cytoplasmic networks is consistent with mechanical integration and entanglement between networks. Based on these extract observations, we propose that MTOCs and astral MTs move away from the midplane as a continuum as they grow after first mitosis, carrying organelles and F-actin with them.

An important question is how well our aster movement systems model centrosome movement in eggs. After anaphase in *Xenopus* eggs, centrosomes move away from the midplane at ~10 μm/min, which is faster than the aster separation movement in Fig 3, and slower than the dynein-based movement over the coverslip in Fig 4. Thus, neither of our extract movement systems precisely reconstituted the speed of aster movement in eggs, but they spanned a wide range of relevant velocities, which increases our confidence that movement is also coherent in eggs. To test whether coherent movement occurs in large eggs, we re-analyzed movies of aster growth and separation movement after first mitosis in live zebrafish eggs expressing a fluorescent MT binding protein (Wühr et al., 2010) (SI Movie 13). Lipid droplets are visible as large dark objects in these movies. These droplets move rapidly and randomly before the aster contacts them, then slowly outwards once they are embedded inside the aster. Using PIV analysis, we observed outward flow of structure in the MT channel at the same speed as the lipid droplets. This analysis suggests large asters in zebrafish eggs also move away from the midzone as a continuum as they grow after first mitosis.

Dynein located throughout the cytoplasm is thought to generate the force that moves asters in large egg cells, but where dynein is anchored has been unclear. Here, we found that all the organelles in the extract moved inwards at the aster surface in a dynein-dependent manner, though this movement was only pronounced when F-actin was fragmented. Thus, all the organelles may serve as dynein anchors, either by recruiting dynein directly, or by physical contact with the ER. The ER moved inwards fastest and to the greatest extent, perhaps because it recruits more dynein, or because individual ER tubules are smaller and more deformable than other organelles. We observed no outward movement of organelles when dynein was inhibited with CC1, so dynein is the dominant microtubule motor in egg asters. The identity of the dynein adapter on egg organelles is unknown. Proteomic analysis estimates ~40 nM HOOK2 and ~10 nM HOOK3 in eggs (Wühr et al., 2014), but preliminary experiments failed to implicate HOOK2 in organelle transport. The only other known dynein adapter present at significant concentration is the kinetochore protein Spindly (SPDL1, ~100 nM) (Wühr et al., 2014). In interphase U2OS cells, Spindly binds to the plasma membrane (Conte et al., 2018). It might be recruited to organelles via its farnesyl modification (Holland et al., 2015) or by interaction with the ZW10-containing NRZ complex (Civril et al., 2010; Menant et al., 2010).

Organelles reproducibly exhibited a burst of inward movement when the growing aster surface first contacted them, then slowed or halted upon incorporation into the aster (Figs 6,7). This burst was much more pronounced when F-actin was fragmented, and it was seen in both mass transport and PIV analyses (Fig 7). Most organelles inside asters were stationary, which explains why the density of organelles in the bulk of the aster is similar to that outside the aster, as previously observed (Hara et al., 2015; Wang et al., 2013). The molecular mechanism that restricts dynein-mediated organelle movement to the aster surface is unknown. F-actin reduced transport speeds of both organelles (Fig 6,7) and artificial cargoes (Fig 8). We hypothesize this is due to drag forces from F-actin networks increasing the effective viscosity of the system for organelles. Organelle speeds remained maximal at the aster surface even when F-actin was fragmented (Fig 7). We hypothesize this is also due to an increase in effective viscosity in the aster interior, due to increased MT density and/or non-motor interactions between organelles and MTs. In contrast, artificial cargoes moved to the aster center at constant speeds whether F-actin was intact or fragmented (Fig 8). Organelles may recruit less dynein than beads, or they may recruit unidentified proteins which act as a brake on dynein-mediated movement (Gurel et al., 2014).

Figure 9 illustrates a working model for aster and centrosome movement in eggs based on the hypothesis that dynein-based forces are restricted to the aster surface. For dynein to pull on astral MTs from organelles, the organelles must move toward the midzone, or at least remain stationary. In contrast, our observations showed that organelles inside asters moved away from the midzone, and inward organelle movement was largely restricted to the aster surface (solid red arrows in Figs 9A,B). Surface force propagates through the aster by mechanical stress, and the aster responds as a deformable gel. Inside asters, MTs act both as a substrate for dynein, as well as a brake on organelles, so dynein may thus generate active stress inside asters, which is different than external force on astral MTs. Asters and centrosomes move apart because the midzone is mechanically softer than the rest of the aster. Continual MT polymerization fills the expanding space at the midzone, and the midzone kinesins Kif4A, Kif20A and Kif23 keep the CPC and centralspindlin focused (Nguyen et al., 2014). Since moving asters advected cytosol (Fig 5), we hypothesize outward translocation of the asters generates hydrodynamic forces that displace cytoplasm around the asters and into the midzone (solid beige arrows in Fig 9C). Actomyosin-based forces make a major contribution to separation movement in our model system (Fig 2), and may also contribute in eggs (hollow green arrows in Fig 9C).

**Figure 9.**
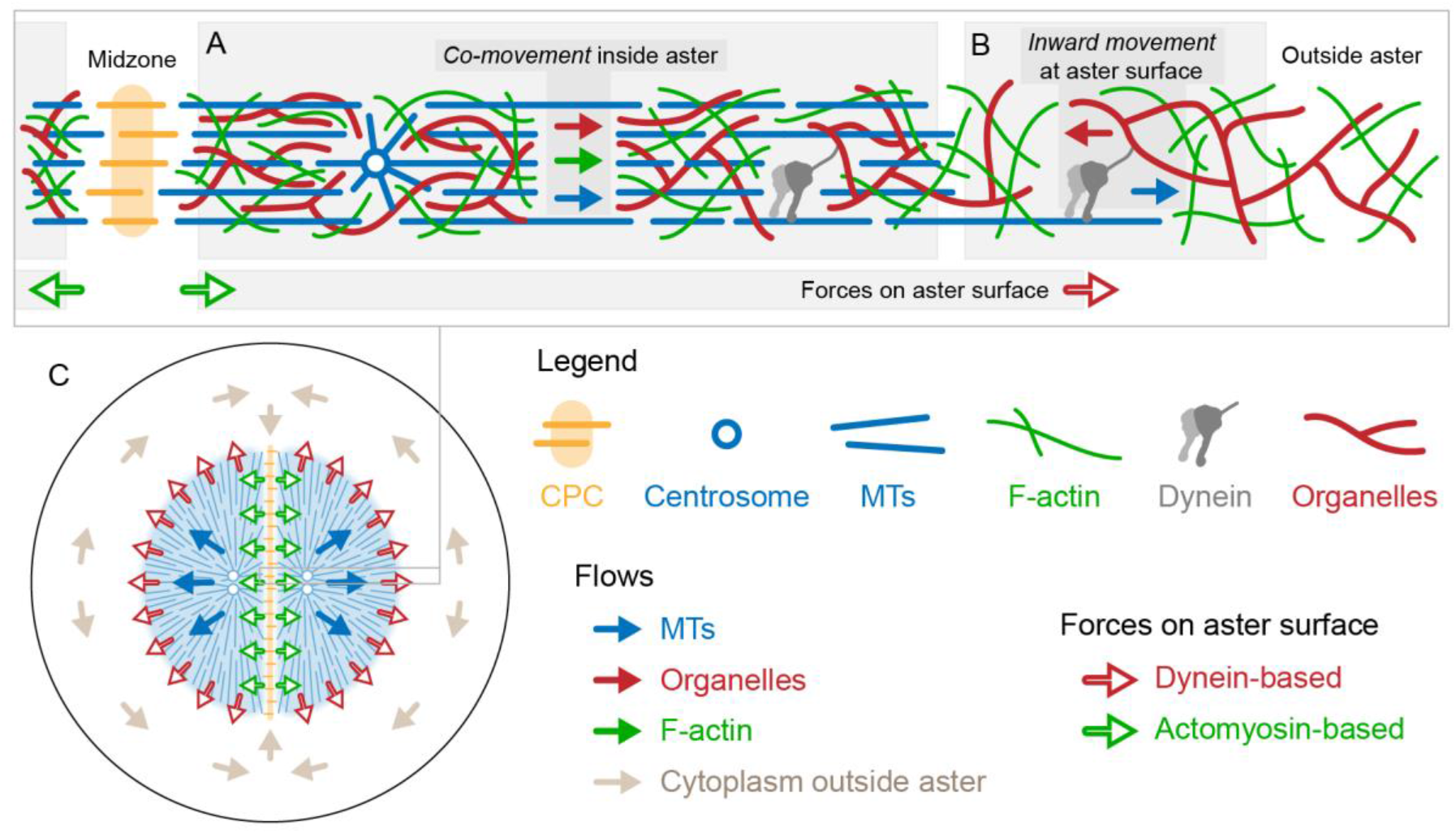
Model of aster positioning based on forces on the aster surface. Solid arrows represent flows of cytoplasmic networks, and hollow arrows represent forces on the aster surface. (A) Inside the aster, cytoplasmic networks move coherently, so there is no dynein pulling force on MTs. (B) At the growing aster surface, organelles move inwards along MTs, generating outward pulling force on MTs (hollow red arrow). Actomyosin contraction away from the midzone, where F-actin is locally disassembled, may also generate forces on asters (hollow green arrows). Surface forces propagate through the aster gel by mechanical stress, causing flow away from the softer midzone. This flow is balanced by net MT polymerization at the midzone. (C) Centrosome separation movement in an egg showing location of forces from dynein and actomyosin and resulting movement of the aster gel and cytoplasm outside the aster. Only the flow of MTs is shown for simplicity, but organelles and F-actin inside asters move with MTs as in panel A.

We have not performed detailed simulations of the model in Fig 9, but conceptually it can account for centrosome separation in an aster pair, as well as centering of a single aster in a spherical space. Frog egg asters exhibit branching MT nucleation, which causes the MT density to remain roughly constant at their surface as the aster grows (Ishihara et al., 2014). If dynein force on the aster surface is proportional to microtubule density, or to organelle density, it will scale linearly with surface area. This “force per unit surface area” feature of the model can potentially account for sperm aster centration in eggs and in regions where MT growth is allowed by UV inactivation of colcemid (Hamaguchi et al., 1986). In extracts, asters are almost 2D and likely experience drag from coverslips, so we expect surface forces may be more significant in 3D eggs than in 2D extracts.

We are not the first to propose aster positioning by surface forces. In early microneedle experiments in echinoderm eggs, Chambers observed that asters behave as a gel and proposed that forces act on their surface (Chambers Jr, 1917). The proposal by Hamaguchi et al. that dynein forces act throughout asters (Hamaguchi et al., 1986) came to dominate recent thinking, but our observations favor the older idea, at least for frog eggs. New experiments are needed to discriminate these models in different systems. The predominance of surface pulling vs length-dependent pulling on asters may depend on the size of the system. We observed dynein-mediated organelle movement relative to MTs over a distance of ~50 μm near the aster surface (Fig 7). This distance corresponds to a relatively thin surface layer in frog egg asters, but it is larger than the cell radius in sea urchin or *C. elegans* eggs. To understand forces on asters in each system, we think it is important to analyze the movement of all cytoplasmic components.

## Materials and methods

### Immunofluorescence

Embryos were fixed and stained as described previously (Field et al., 2019). Embryos were fixed in 90% methanol, 10% water, 50 mM EGTA pH 6.8 for 24 h at room temperature with gentle shaking. After fixation, embryos were rehydrated in steps from 75%, 50%, 25%, to 0% methanol in TBS (50 mM Tris pH 7.5, 150 mM NaCl) for 15 min each step with gentle shaking. Rehydrated embryos in TBS were cut in half on an agarose cushion using a small razor blade. Before staining, embryos were bleached overnight in 1% hydrogen peroxide, 5% formamide (Sigma-Aldrich #F9037), 0.5x SSC (75 mM NaCl, 8 mM sodium citrate pH 7). To stain, embryos were incubated with directly labeled antibodies at 0.5-2 μg/mL for at least 24 h at 4 °C with very gentle rotation. Antibodies were diluted in TBSN (10 mM Tris-Cl pH 7.4, 155 mM NaCl, 1% IGEPAL CA-630 (Sigma-Aldrich #I8896), 1% BSA, 2% FCS, 0.1% sodium azide). After antibody incubation, embryos were washed in TBSN for at least 48 h with several solution changes, then washed once in TBS and twice in methanol, with methanol washes for 40 min each. Embryos were cleared in Murray clear solution (benzyl benzoate (Sigma-Aldrich #B6630)/benzyl alcohol (Sigma-Aldrich #402834) 2:1). Embryos were mounted in metal slides 1.2 mm thick with a hole in the center. The hole was closed by sealing a coverslip to the bottom of the slide using heated Parafilm.

Endoplasmic reticulum (ER) was labeled with an anti-LNPK antibody (Wang et al., 2016) directly labeled with Alexa Fluor 568 NHS ester (Thermo Fisher #A20003). The ER was also probed with labeled anti-Protein disulfide-isomerase A3 (PDIA3) (Boster Bio #PB9772). PDIA3 is an ER lumen protein and had a similar distribution as the anti-LNPK antibody (not shown). Microtubules were labeled with an anti-tubulin antibody directly labeled with Alexa Fluor 647 NHS ester (Thermo Fisher #A20106).

### Extract preparations

Actin-intact, CSF *Xenopus* egg extract was prepared as described previously (Field et al., 2017). CSF extracts were stored at 4-10 °C and flicked occasionally to disperse membranes. Extracts stored in this way were typically usable for ~8 h. Before each reaction, extracts were cooled on ice to ensure depolymerization of cytoskeletal networks.

### Interphase aster assembly reactions

In a typical reaction, fluorescent probes were added to CSF extract on ice. To trigger exit from CSF arrest and entry to interphase, calcium chloride was added to 0.4 mM final concentration. To ensure complete progression to interphase, the reaction was mixed well immediately after calcium addition by gently flicking and pipetting. Extracts were pipetted using 200 μL pipette tips manually cut to a wider bore to reduce shear damage, which can make membranes in the extract appear coarser by eye. Reactions were incubated in an 18 °C water bath for 5 min then returned to ice for 3 min. Next, drugs or dominant negative constructs were added (see Perturbations below), and in some cases reactions were split for direct comparison between control and perturbed conditions. Last, Dynabeads Protein G (Thermo Fisher #10004D) functionalized with an activating anti-Aurora kinase A (anti-AURKA) antibody were added as artificial microtubule organizing centers (MTOCs) (Tsai et al., 2005). For experiments in which asters moved away from one another, unlabeled anti-INCENP antibody was included at a final concentration of 4 nM to promote zone formation by activating the CPC.

### Coverslip passivation

18 and 22 mm square coverslips were passivated with poly-L-lysine covalently grafted to polyethylene glycol (PLL-g-PEG) (SuSoS #PLL(20)-g3.5-PEG(2)) as described previously (Field et al., 2019). Coverslips were cleaned by dipping them in 70% ethanol, igniting the ethanol with a gas burner, cooling the coverslips for several seconds, then the coverslips were passivated by placing them on a droplet of 0.1 mg/mL PLL-g-PEG in 10 mM HEPES pH 7.4 on Parafilm. 18 mm coverslips were placed on 90 μL droplets, and 22 mm coverslips were placed on 110 μL droplets. After 30 min incubation, excess PLL-g-PEG was rinsed by placing coverslips on droplets of distilled water twice for 5 min each, then drying them with a stream of nitrogen gas. To check the passivation, when we focused near the coverslips, we found no evidence of a surface layer of cytoskeletal filaments or organelles adsorbed to the coverslips. Quite the opposite, the density of cytoplasmic networks was typically lower near the coverslips and higher near the midplane between the coverslips, we suspect due to continuous contraction of actomyosin away from the coverslips sustained by continuous diffusion of monomer toward the coverslips.

### Flow cell assembly

Flow cells were assembled from the passivated coverslips to increase physical stability of the system and reduce global flows. 22 mm square coverslips were sealed to a metal slide holder via a thin layer of molten VALAP (Vaseline, lanolin (Sigma-Aldrich #L7387), paraffin (Sigma-Aldrich # 327204) 1:1:1 by mass). Then an 18 mm square coverslip was immobilized above the 22 mm coverslip using two pieces of thin double-sided tape (3M Extended Liner Tape #920XL) spaced ~1 cm apart. The tape has a nominal thickness of 25 μm and resulted in flow cells ~20 μm deep after pressing the coverslips together.

### Imaging

Extract reactions were perfused into flow cells, then the edges were sealed with VALAP. In experiments with a single condition, imaging was started immediately. In experiments with multiple conditions imaged in parallel, the slide holder was first chilled on ice for several seconds, so aster growth would start at the same time across conditions. Extracts were imaged on a Nikon Eclipse Ti2-E inverted microscope with Nikon CFI Plan Apo Lambda 20x, NA 0.75 objective lens, SOLA SE V-nIR light engine, and with either a Nikon DS-Qi2 or Andor Zyla 4.2 PLUS sCMOS camera. The microscope room was cooled to less than 20 °C, otherwise spontaneously nucleated microtubules can overtake reactions. Throughout the paper, time is measured with respect to warming the reaction and the start of aster growth. Depending on the MTOC density, asters typically grew into contact at 8-15 min and formed CPC-positive interaction zones several minutes later.

### Fluorescent probes

Microtubules (MTs) were imaged with either bovine or frog tubulin directly labeled with Alexa Fluor 647 at a final concentration of 250 nM, or with a phosphodeficient version of the microtubule binding domain of Tau fused to mCherry (Mooney et al., 2017) at a final concentration of 20 nM. Growing +TIPs of MTs were labeled with EB1-GFP at a final concentration of 110 nM. The chromosomal passenger complex (CPC) was labeled with an anti-INCENP antibody directly labeled with Alexa Fluor 647 at a final concentration of 4 nM. Endoplasmic reticulum (ER) was labeled with DiI (1,1’-dioctadecyl-3,3,3’,3’-tetramethylindocarbocyanine perchlorate, aka DiIC18(3)) (Thermo Fisher #D282) or DiD (1,1’-dioctadecyl-3,3,3’,3’-tetramethylindodicarbocyanine perchlorate, aka DiIC18(5)) (Thermo Fisher #D307) at a final concentration of 4 μg/mL. Mitochondria were labeled with tetramethylrhodamine, ethyl ester (TMRE) (Thermo Fisher #T669) at a final concentration of 0.3 μg/mL. Acidic organelles were labeled with LysoTracker Red DND-99 (Thermo Fisher #L7528) at a final concentration of 130 nM. To allow these dyes to pre-incorporate into the membranous organelles, especially important for the DiI and DiD, stock solutions were first dissolved in DMSO to a concentration of 2 mg/mL (DiI/DiD), 0.2 mg/mL (TMRE), or 200 μM (LysoTracker Red), then diluted 50 fold into extract. These extract working solutions were incubated in an 18 °C water bath for 45 min, flicking every 15 min to disperse membranes. Then the extract working solutions were stored on ice until use, then diluted an additional 10-30 fold into the final reaction. F-actin was imaged with Lifeact-GFP (Moorhouse et al., 2015; Riedl et al., 2008) at a final concentration of 300 nM. More details on fluorescent probes are reported in (Field et al., 2017).

### Perturbations

To fragment F-actin, Cytochalasin D (CytoD) was added to a final concentration of 20 μg/mL. CytoD was diluted in DMSO to 10 mg/mL, then diluted 20 fold into extract. This extract working solution was stored on ice until use, then diluted an additional 25 fold into the final reaction. CytoD and other drugs or dominant negative constructs were typically added to actin-intact extracts after cycling to interphase, then reactions were split for direct comparison between control and perturbed extracts. Alternatively, CytoD may be added during extract preparations before the crushing spin, following the classic CSF extract protocol (Murray, 1991). The ER appeared coarser in CytoD extracts than in actin-intact extracts, and the ER appeared to coarsen over time in actin-intact extracts plus CytoD.

To inhibit dynein, the p150-CC1 fragment of dynactin (King et al., 2003) acts as a dominant negative for dynein function was added to a final concentration of 40 μg/mL.

### HOOK2 antibody

An affinity purified C-terminal peptide antibody was produced in rabbit against *Xenopus laevis* HOOK2 (C-SRSHTLLPRYTDKRQSLS) (Cocalico Biologicals, Inc., PA).

### HOOK2 immunoprecipitation-mass spectrometry (IP-MS)

Dynabeads Protein G (Thermo Fisher #10004D) (20 μL Dynabeads slurry per reaction) were saturated with rabbit IgG (anti-HOOK2 or random IgG (Jackson ImmunoResearch #011-000-003)) by overnight binding, then washed 3x with CSF-XB (100 mM KCl, 2 mM MgCl2, 0.1 mM CaCl2, 10 mM K HEPES pH 7.7, 5 mM EGTA, 50 mM sucrose). Each immunoprecipitation reaction contained 150 μL interphase or CSF-arrested egg extract treated with 10 μg/mL Cytochalasin D to inhibit gelation. Extract plus Dynabeads was rotated gently for 60 min at 4 °C, then washed 4x in 50 mM KCl, 1 mM MgCl2, 10 mM K HEPES pH 7.7, 1 mM EGTA at 0 °C. The tubes were changed twice during the washes to remove extract protein bound to their walls. Protein bound to the Dynabeads was eluted in 20 μL of 5 M guanidine thiocyanate, 5 mM dithiothreitol (DTT) (US Biological #D8070) for 10 min at 60 °C, then cysteines were alkylated with N-ethylmaleimide (NEM). The eluate was precipitated with chloroform-methanol then subjected to proteolysis followed by TMT labeling as described (Sonnett et al., 2018).

### Preparation of dynein on coverslips

Coverslips were passivated following the protocol above but using biotinylated PLL-g-PEG. NeutrAvidin and biotinylated Protein A (GenScript #M00095) were mixed in a 1:1 ratio to a final concentration of 10 μM and stored at 4 °C. Just before functionalizing the coverslips, the NeutrAvidin (Thermo Fisher #31000) and biotinylated Protein A mixture (GenScript #M00095) was diluted 42 fold to 240 nM in 1x PBS with 0.0025% Tween 20. That concentration was found to be the smallest amount to decrease the surface tension enough to maintain a layer of solution on the coverslips, to reduce damage to the functionalized surfaces due to air-water interfaces when transferring the coverslips from one droplet to another. Coverslips were incubated with the NeutrAvidin and biotinylated protein A mixture at least 30 min on droplets on Parafilm at room temperature. Coverslips were incubated under a box with a damp paper towel, to block room light and to reduce evaporation. After the incubation, coverslips were rinsed twice on droplets of 1x PBS with 0.0025% Tween 20 for 5 min each, then incubated with anti-HOOK2 or random IgG diluted in 1x PBS with 0.0025% to a final concentration of 10 μg/mL at least 30 min. After the incubation with antibody, coverslips were rinsed twice on droplets of 1x PBS with 0.0025% Tween 20, then twice on droplets of distilled water, then swirled in a beaker of distilled water, then gently drying them with a stream of nitrogen gas. Coverslips were often used same day, but could be stored overnight in the dark at 4 °C and used the following day. After perfusing extracts into flow cells and sealing the edges with VALAP, the metal slide holders were chilled for 10 min on a metal block on ice, to allow endogenous HOOK2 and dynein-dynactin time to bind the anti-HOOK2 before the start of aster growth.

### Photo-release of fluorescein from MTOCs

Caged fluorescein with -O-CH2-COOH functionality on the caging groups was synthesized as described (Mitchison et al., 1998). Carboxylic acid groups were activated as sulfo-NHS esters in a small reaction containing 2 micromols caged fluorescein, 5 micromols sodium sulfo-NHS and 5 micromols 1-ethyl-3-(3-dimethylaminopropyl)carbodiimide (EDC) (Thermo Fisher #22980) in 10 μL of DMSO. After 1 h at room temperature, this reaction mix was added directly to protein coated beads. Direct modification of anti-AURKA beads caused loss of nucleation activity, so we first biotinylated beads, then modified with caged fluorescein, then attached anti-AURKA IgG using a NeutrAvidin bridge. Dynabeads Protein A (Thermo Fisher #10002D) were sequentially incubated with goat anti-rabbit whole serum (Jackson ImmunoResearch #111-001-001) then biotinylated rabbit IgG (homemade). They were labeled with the caged fluorescein reaction mix in 0.1 M K HEPES pH 7.7 for 1 h, then washed again. We empirically titrated the amount of reaction mix added such that beads were maximally labeled while still retaining nucleation activity in extract. After labeling with caged fluorescein, beads were incubated sequentially with a mixture of NeutrAvidin and biotinylated protein A, then rabbit anti-AURKA to confer nucleation activity. Pure proteins were added at 10-20 μg/ml and serum was added at 1/20. All binding reactions were incubated for 20 min, and washes were in 1x PBS. Fluorescein was released from beads by exposing the microscope field to full illumination in the DAPI channel (395 nm) for 5 sec.

## Analyses

### PIV

PIVlab (Thielicke et al., 2014) was used to estimate flow fields of cytoplasmic networks based on particle image velocimetry (PIV). Though PIV is primarily used to estimate flow fields based on tracer particles embedded in fluids, PIV has been used to estimate cortical or cytoplasmic flows in *C. elegans* cortices (Mayer et al., 2010), zebrafish epithelia (Behrndt et al., 2012), and *Drosophila* embryos (Deneke et al., 2019). Likewise, cytoplasmic networks in the *Xenopus* egg extracts included structures with sufficient contrast for PIV. The cytoplasmic networks exhibited dynamic turnover, so it was important to image with a time interval short enough to retain sufficient correlation between frames for PIV. For example, the time scale for F-actin turnover was ~1 min, based on recovery of F-actin in a region where F-actin had been mechanically cleared, consistent with estimates based on measurements of network density and flow in contractile actomyosin networks (Malik-Garbi et al., 2019). Time intervals less than 20 s worked well for PIV.

### Gaussian fitting of photo-released fluorescein

2D Gaussian fitting of fluorescein photo-released from MTOCs was performed using a nonlinear least squares solver in MATLAB (Nootz, 2020). After photo-release the MTOCs were bright due to uncaged fluorescein that remained bound to the MTOCs. Thus the MTOCs were masked as not to bias the Gaussian fits. We fit expansion of the fluorescein cloud to a model of diffusion, and we assumed a diffusion coefficient of fluorescein in water of 425 μm^2^/s (Culbertson et al., 2002). Advection of cytosol with cytoplasmic networks is consistent with a poroelastic Péclet number *VLμ*/*Eξ*^2^ greater than unity (Mitchison et al., 2008; Moeendarbary et al., 2013). Given the oscillatory speed *V* ~1 μm/s (Fig 4D) and amplitude *L* ~30 μm (Fig 4C), and assuming a viscosity *μ* ~6x water (Fig 5E) and an elastic modulus *E* ~10 Pa (Valentine et al., 2005), we estimate the upper bound on the effective pore size *ξ* of cytoplasmic networks in this system is ~100 nm.

### Analysis of organelle mass transport

The flux-based analysis of organelle transport is described in Fig S4. In summary, images were background subtracted and flat field corrected, then a region of interest (ROI) was defined large enough to enclose the aster at all time points, so the total amount of ER in the ROI was conserved. Then, the total intensity was normalized across frames to correct for photobleaching. The net flux of organelle fluorescence intensity toward MTOCs was calculated as described in Fig S4. In particular, the average intensity was calculated in annular bins with a width of 10 μm, then the cumulative total intensity was calculated from the MTOC to outside the aster, then the net flux was calculated at each radial distance by subtracting subsequent cumulative total intensity profiles.

## Acknowledgements

This work was supported by NIH grant R35GM131753 (TJM) and MBL fellowships from the Evans Foundation, MBL Associates, and the Colwin Fund (TJM and CMF). JFP was supported by the Fannie and John Hertz Foundation, the Fakhri lab at MIT, the MIT Department of Physics, and the MIT Center for Bits and Atoms. The authors thank the Nikon Imaging Center at Harvard Medical School and Nikon at MBL for imaging support, and the National Xenopus Resource at MBL for support. The authors thank Keisuke Ishihara and Luolan Bai for critical feedback on the manuscript, and thank Martin Wühr, Jay Gatlin, Nikta Fakhri, David Burgess, Fabian Romano-Chernac, Sam Reck-Peterson, and Mark Terasaki for helpful conversations. The authors thank Martin Wühr for the movie showing post-anaphase separation movement of asters after first mitosis in zebrafish. The anti-LNPK antibody was a gift from Tom Rapoport (Harvard Medical School and Howard Hughes Medical Institute, Boston, MA). The Tau-mCherry construct was a gift from Jay Gatlin (University of Wyoming, Laramie, WY). The EB1-GFP construct was a gift from Kevin Slep (UNC Chapel Hill, NC). The Lifeact-GFP construct was a gift from David Burgess (Boston College, Newton, MA).

## Author contributions

JFP, CMF, and TJM designed and conducted the experiments, analyzed the data, and wrote the manuscript. SF helped with the analyses in Figs 3D, 4F, and 7C,F,I,J, and read and edited the manuscript. MS helped with sample preparations and performed mass spectrometry (MS) measurements for Fig S2.

## Supporting information

**SI Figure 1.**
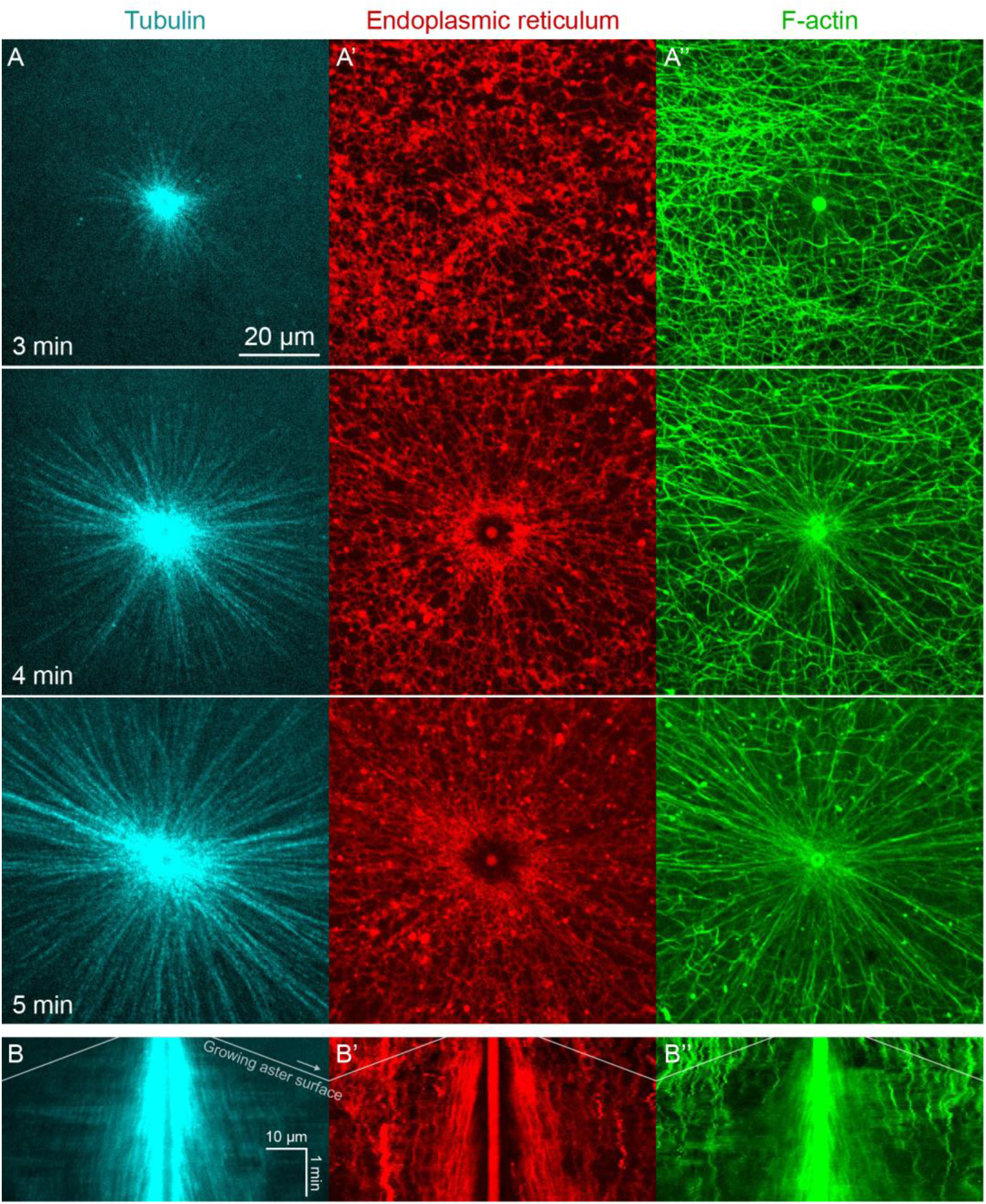
Growth of astral MTs from MTOCs, and dynamic reorganization of MTs, ER, and F-actin. (Related to Fig 1) (A) MTs imaged at near-speckle concentrations with tubulin-Alexa Fluor 647, ER labeled with DiI, and F-actin labeled with Lifeact-GFP. Images taken on a spinning disk confocal microscope with 60x objective (see SI Movie 1). (B) Kymographs generated along a line through the MTOC. The white diagonal lines indicate the growing aster surface.

**SI Figure 2.**
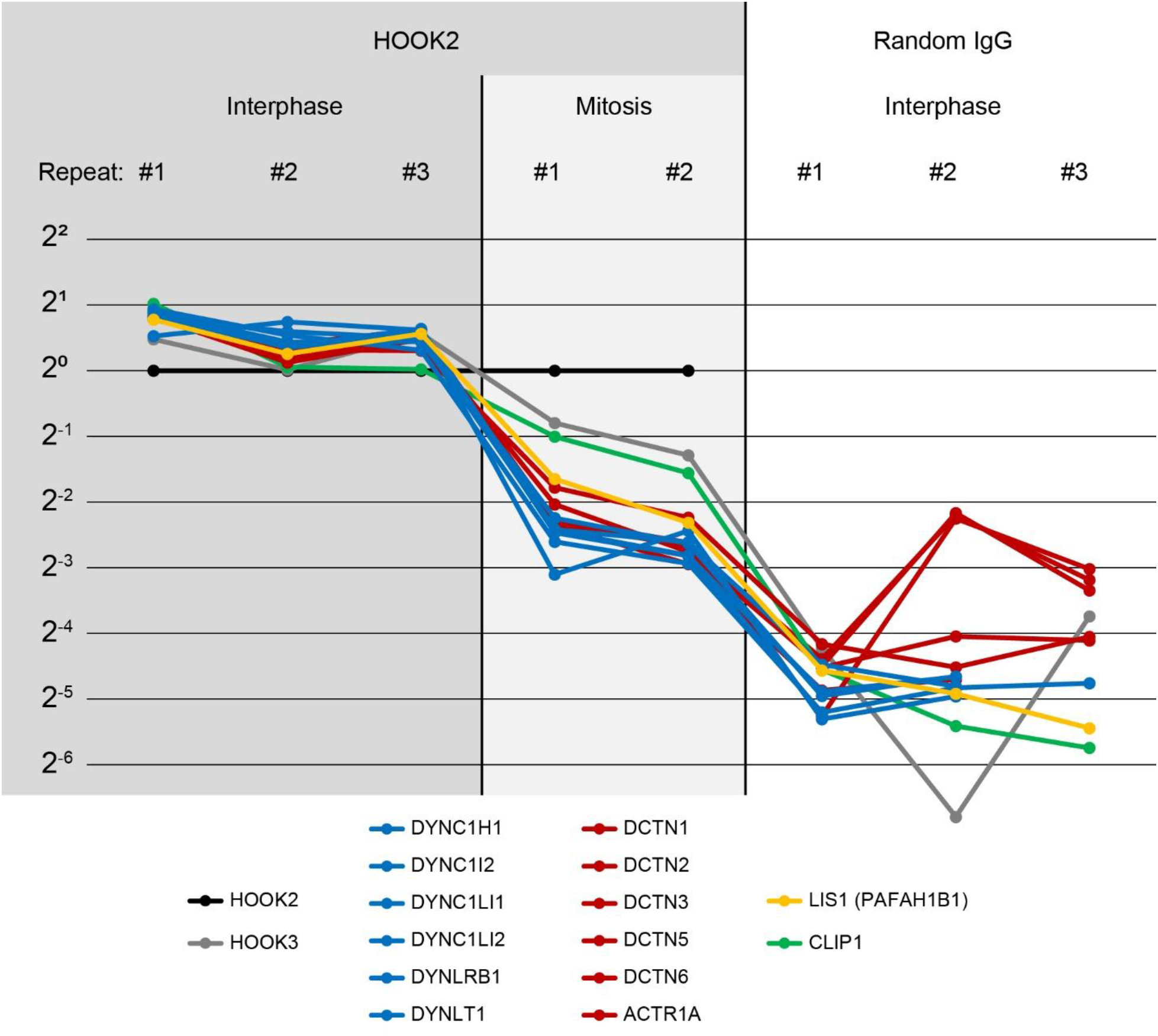
Characterization of the HOOK2 C-terminal peptide antibody. (Related to Figures 4, 5, and 8) We performed immunoprecipitation-mass spectrometry (IP-MS) on Protein G Dynabeads functionalized with anti-HOOK2. In 3 repeat extracts, we measured 3 conditions: anti-HOOK2 in interphase extracts (columns 1 to 3), anti-HOOK2 in mitotic extracts (columns 4 and 5), and as negative control, random IgG in interphase (columns 6 through 8). HOOK2 conditions were normalized so the amount of HOOK2 was constant. Random IgG conditions were normalized to have the same IgG count as the average IgG count of the HOOK2 columns. Then for each protein, the sum across channels was normalized, so the abundance values represent relative enrichment across the 8 channels. Abundances are shown on a log scale with base 2. We here show dynein subunits (blue), dynactin subunits (red), and other proteins known to interact with dynein-dynactin (yellow, green) that came down in the immunoprecipitations. The interaction between HOOK2 and the dynein-dynactin complex was stronger in interphase (left columns) than in mitosis (middle columns).

**SI Figure 3.**
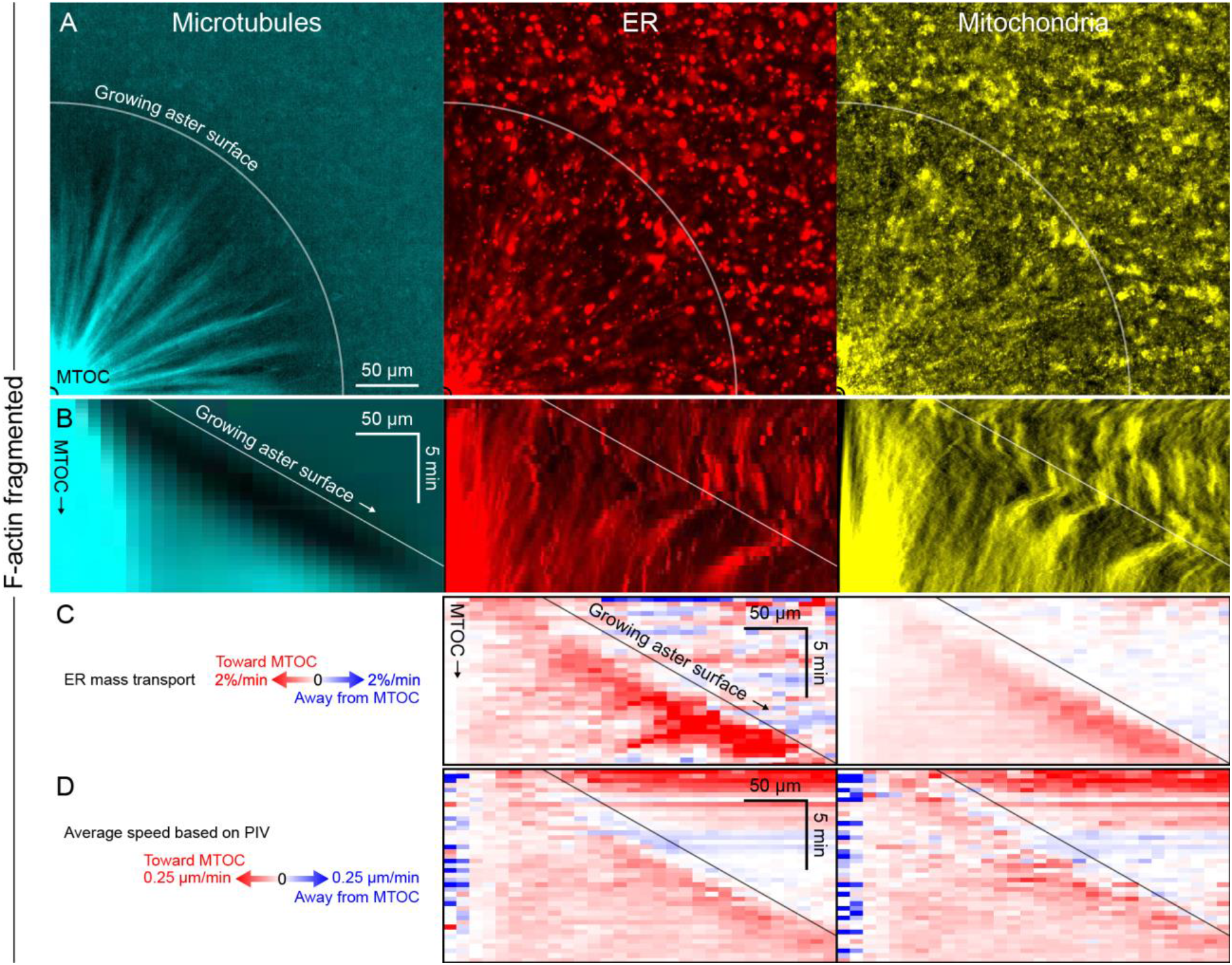
Like other organelles, mitochondria exhibited a burst of organelle movement near the growing aster surface. (Related to Figs 6 and 7) (A) F-actin was fragmented by Cytochalasin D. Microtubules were labeled with tubulin-Alexa Fluor 647, ER with DiI, and mitochondria with TMRE (see SI Movie 10). (B) Kymographs along a line from the MTOC. (C) Mass transport map for ER and mitochondria averaged over a quadrant, in the same frame of reference as the kymographs in panel B. Mass transport analysis is described in Fig S4. (D) Average speed based on PIV, in the same frame of reference as panels B and C, and averaged over a quadrant. In all analysis methods, mitochondria behaved similar to acidic organelles and likewise exhibited a burst of movement at the growing aster surface.

**SI Figure 4.**
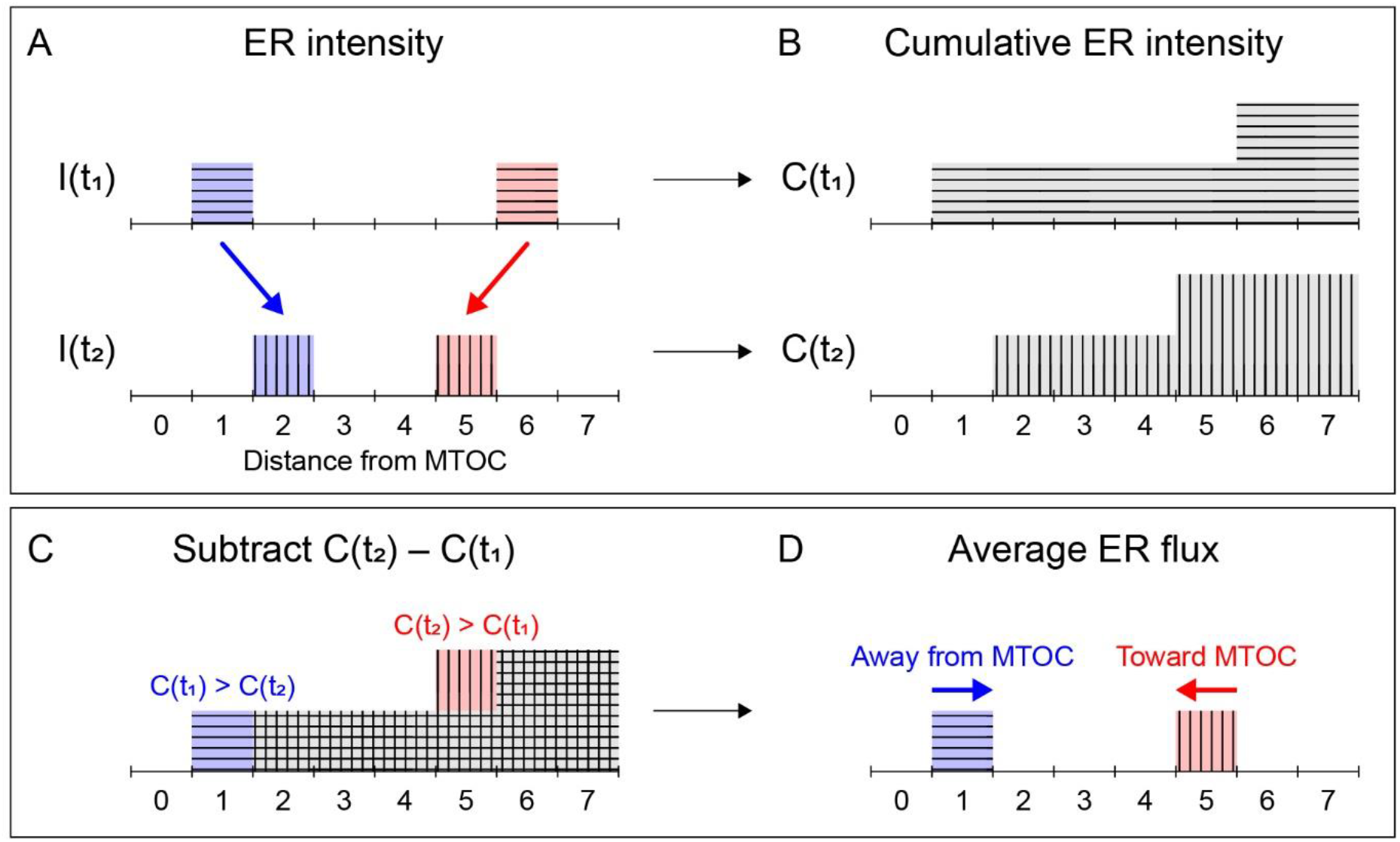
Explanation of flux-based analysis of organelle transport. (Related to Fig 7) (A) From the first frame (top row, horizontal bars) to the second frame (bottom row, vertical bars), consider one unit of ER that moves from left to right (blue), and another unit of ER that moves from right to left (red). (B) Cumulative intensity of ER during the first frame (horizontal bars) and second frame (vertical bars). (C) Difference between the cumulative intensity during the second frame C(t2) (vertical bars) minus the cumulative intensity during the first frame C(t1) (horizontal bars) highlights mass transport of the ER toward the MTOC. (D) There is a net flux of ER toward the MTOC where C(t2) exceeds C(t1) (red, vertical bars), while there is a net flux of ER away from the MTOC where C(t1) exceeds C(t2) (blue, horizontal bars).

**SI Figure 5.**
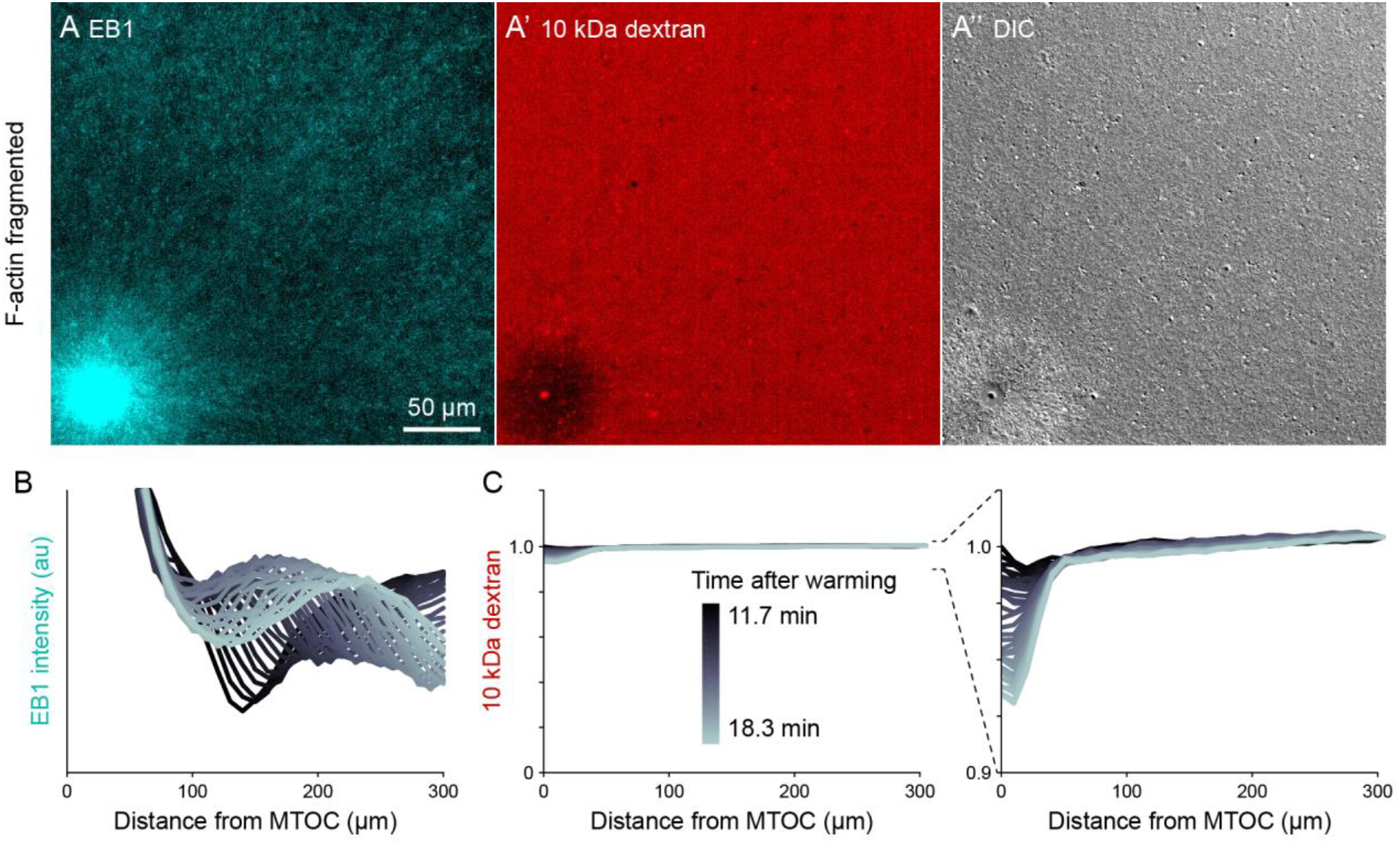
Dextran was excluded in organelle-rich region within ~50 μm of MTOCs. (Related to Figs 7 and 8) (A) The growing +TIPs of MTs were labeled with EB1-GFP, the relative concentration of cytosol was inferred from 10 kDa dextran labeled with Alexa-Fluor 568, and organelles were visualized with differential interference contrast (DIC). Images taken 18.3 min after warming to 20 °C and the onset of aster growth. (B,C) Intensity profiles away from the MTOC colored by time point. In the EB1 graph, the wavefront corresponds to the growing aster surface. To correct for photobleaching, dextran profiles were normalized to the intensity at the point farthest from the MTOC.

### SI Movies

**SI Movie 1. Dynamic reorganization of cytoplasmic networks during the initial stages of aster nucleation and growth.** Related to Fig 1. MTs were labeled with tubulin-Alexa Fluor 647, ER with DiI, and F-actin with Lifeact-GFP. Imaged on a spinning disk confocal with 60x objective lens. Cytoplasmic networks were highly dynamic, and astral MTs dynamically reorganized the ER and F-actin networks. Parts of the ER exhibited abrupt and transient motion toward the MTOC, presumably driven by dynein, and the F-actin transitioned from random to radial entrainment with MTs.

**SI Movie 2. Co-movement of MTs, ER, and F-actin during aster separation movement.** Related to Fig 2A and Fig 3. MTs were labeled with tubulin-Alexa Fluor 647, ER with DiI, F-actin with Lifeact-GFP, and organelles were shown in differential interference contrast (DIC). All cytoplasmic networks moved together. Note the flow of organelles visible in DIC: inside asters, where the density of F-actin, MTs, and ER was higher, organelles flowed with the asters; whereas along interaction zones between asters where the density of F-actin was lower, organelles flowed in the opposite direction, into the space on the right that was vacated by the asters moving to the left.

**SI Movie 3. Both dynein and actomyosin contribute to aster separation movement.** Related to Fig 2. We compared four conditions: control with F-actin intact, dynein inhibited by CC1, F-actin fragmented by Cytochalasin D, and double inhibition of dynein and F-actin. F-actin was labeled with Lifeact-GFP, ER with DiI, and CPC-positive interaction zones with anti-INCENP-Alexa Fluor 647. MTs grew and CPC-positive zones formed between asters in all conditions. F-actin and ER were imaged instead of MTs because local disassembly of F-actin along CPC-positive interaction zones enables aster separation movement, and inward transport of ER and other organelles is thought to drive dynein-based aster movement.

**SI Movie 4. Co-movement of MTs, ER, and F-actin during oscillatory aster movement on coverslips functionalized with dynein.** Related to Fig 4. MTs were labeled with tubulin-Alexa Fluor 647, ER with DiI, and F-actin with Lifeact-GFP. All cytoplasmic networks moved together. Dynein was recruited to coverslips via an antibody to the endogenous dynein adapter HOOK2.

**SI Movie 5. Co-movement of keratin with moving asters during oscillatory aster movement.** Related to Fig 4. MTs were labeled with tubulin-Alexa Fluor 647, F-actin with Lifeact-GFP, and keratin with anti-keratin-Alexa Fluor 568. All cytoplasmic networks moved together.

**SI Movie 6. Advection of fluorescein with moving asters during oscillatory aster movement.** Related to Fig 5. The first frames show MTs labeled with tubulin-Alexa Fluor 647, and the aster filled the region. The next few frames show the caged fluorescein attached to the MTOC. Then, the fluorescein was simultaneously photo-released from the MTOC as its fluorescence was uncaged, releasing a cloud of fluorescent fluorescein around the MTOC. The fluorescein cloud was fit with a 2D Gaussian. The center of the cloud is indicated at the intersection of the red and green lines, and the standard deviation of the cloud is indicated by the black circle. The plots above and to the right indicate the intensity values along the lines, and the black curves show the 2D Gaussian fit along the lines.

**SI Movie 7. F-actin reduced dynein-based transport of ER and mitochondria on stationary asters.** Related to Fig 6. The growing aster is indicated by growing +TIPs labeled with EB1-GFP, ER was labeled with DiI, and mitochondria with TMRE. In control with intact F-actin, some ER accumulated around the MTOC, and little to no mitochondria accumulated around the MTOC. When F-actin was fragmented, a greater fraction of ER and mitochondria were transported toward the MTOC. When dynein was inhibited, organelles were not transported, neither toward nor away from the MTOC.

**SI Movie 8. Burst of ER movement at the growing aster surface in control with F-actin intact.** Related to Fig 7. MTs were labeled with tubulin-Alexa Fluor 647, ER with DiI, and F-actin with Lifeact-GFP. The ER exhibited a burst of movement toward the MTOC at the growing aster surface, resulting in transient depletion of the ER intensity near the aster surface.

**SI Movie 9. Burst of ER and acidic organelle movement at the growing aster surface with F-actin fragmented.** Related to Fig 7. Transport of ER and acidic organelles with F-actin fragmented by Cytochalasin D. MTs were labeled with tubulin-Alexa Fluor 647, ER with DiI, and acidic organelles with LysoTracker Red. Unlike in control with F-actin intact, the burst of movement near the aster surface was highly reproducible when F-actin was fragmented with Cytochalasin D.

**SI Movie 10. Burst of ER and mitochondria movement at the growing aster surface with F-actin fragmented.** Related to Fig S3. Same experiment as SI Movie 9. Mitochondria were labeled with TMRE.

**SI Movie 11. Artificial cargoes, Dynabeads functionalized with dynein via anti-HOOK2, were transported at constant speeds throughout asters.** Related to Fig 8. MTs were labeled with tubulin-Alexa Fluor 488, anti-HOOK2 beads with Fab fragment-Alexa Fluor 568, and negative control beads were functionalized with random rabbit IgG and labeled with Fab fragment-Alexa Fluor 647.

**SI Movie 12. Exclusion of 10 kDa dextran from the organelle-rich region around MTOCs.** Related to Fig S5. 10 kDa dextran labeled with Alexa Fluor 568 was excluded in a ~50 μm radius around the MTOC.

**SI Movie 13. Post-anaphase aster separation movement in a zebrafish embryo consistent with continuum movement.** Related to Fig 9. Movie from (Wühr et al., 2010) and analyzed with permission. Microtubules were labeled with microtubule-binding domain of Ensconsin fused to three GFPs (EMTB-3GFP) (Faire et al., 1999; von Dassow et al., 2009). Flows of MTs were estimated by PIV (Methods).

